# Combining transgenesis with paratransgenesis to fight malaria

**DOI:** 10.1101/2022.03.02.482642

**Authors:** Wei Huang, Joel Vega-Rodriguez, Christopher Kizito, Sung-Jae Cha, Marcelo Jacobs-Lorena

## Abstract

Malaria is among the deadliest infectious diseases and *Plasmodium*, the causative agent, needs to complete a complex development cycle in its vector mosquito for transmission to occur. Two promising strategies to curb transmission are transgenesis, consisting of genetically engineering mosquitoes to express anti-malarial effector molecules and paratransgenesis, consisting of introducing into the mosquito, commensal bacteria engineered to express anti-malarial effector molecules. Although both approaches restrict parasite development in the mosquito, it is not known how their effectiveness compares. Here we provide an in-depth assessment of transgenesis and paratransgenesis and evaluate the combination of the two approaches. Using the Q-system to drive gene expression, we engineered mosquitoes to produce and secrete two effectors – scorpine and the MP2 peptide – into the mosquito gut and salivary glands. We also engineered *Serratia*, a commensal bacterium capable to spread through mosquito populations, to secrete the same two effectors into the mosquito gut. Whereas both mosquito-based and bacteria-based approaches strongly reduced the oocyst and sporozoite intensity, a substantially stronger reduction of *P. falciparum* development was achieved when transgenesis and paratransgenesis were combined. Most importantly, transmission of *P. berghei* from infected to naïve mice was maximally inhibited by the combination of the two approaches. Combining these two strategies promise to become a powerful approach to combat malaria.

**Significance:** Malaria kills hundreds of thousand persons yearly. Clearly, new approaches are needed to fight this disease. Two promising approaches are based on the concept of genetically modifying the mosquito to make it a poor vector for the parasite: 1) transgenesis (engineering the mosquito to deliver anti-malarial compounds) and 2) paratransgenesis (engineering mosquito symbiotic bacteria to deliver anti-malarial compounds). The key questions addressed by this manuscript are: which of the two is the most promising approach? And because transgenesis and paratransgenesis are not mutually exclusive, would the combination of both be the most effective strategy? Our results argue for the combination of the two, showing the additive impact that these two strategies may have in controlling malaria transmission in the field.

## Introduction

Over 400,000 people, mostly young African children, died of malaria in 2019 (1). Whereas world malaria incidence has declined by 27% during the first 15 years of this century, in the last four years it declined by less than 2%, indicating that current interventions to control this deadly disease are waning.^1^ The development of innovative approaches to reduce this intolerable burden is sorely needed.

The strategy of targeting the mosquito to fight malaria is based on two premises: 1) the mosquito is an obligatory vector for parasite transmission and 2) strong bottlenecks limit parasite development in the mosquito and during transmission to the mammalian host.^2^ The mosquito acquires the parasite when it bites an infected individual. Of the large numbers of gametocytes (∼10^4^) ingested by the mosquito, only a few (single digits) ookinetes succeed in traversing the mosquito gut and differentiate into oocysts, defining the first strong bottleneck.^3^ Each oocyst produces thousands of sporozoites, a good proportion of which invade the salivary glands, where they are stored. Only a small number of these sporozoites (on the order of 1% of total salivary gland content) are delivered when an infected mosquito bites a new individual, defining a second strong bottleneck. ^4^

Since the early demonstration that mosquitoes can be engineered to be refractory to the parasite,^5^ the effectiveness of this approach has been robustly demonstrated in the laboratory by simultaneous expression of multiple effector genes (genes that stop parasite development without affecting the mosquito vector).^6, 7^ The major current challenge is to devise means to introduce the genes that confer refractoriness into mosquito populations. This will most likely be achieved by use of CRISPR/Cas9 gene drives.^8, 9^ In addition to technical aspects, topics to be resolved include regulatory and ethical issues related to the release of genetically modified organisms in nature.

An independent approach to suppress the mosquito vectorial capacity is to express effector genes from symbiotic bacteria rather than from the mosquito itself, an approach referred to as paratransgenesis. Paratransgenesis has the advantage that the bacteria occur in the mosquito gut in large numbers, in close proximity to the most vulnerable parasite forms. Since the early demonstration of the effectiveness of paratransgenesis to contain the spread of *Trypanosoma cruzi*, the causative agent of Chagas disease, by the *Rhodnius prolixus* vector,^10^ this approach has been developed for suppressing the mosquito’s ability to vector the malaria parasite.^6, 11, 12, 13^ As is the case for gene drive, the mosquito symbiont *Serratia AS1* can spread through mosquito populations and be engineered to secrete effector proteins.^14^

This work addresses two unanswered questions: 1) which of the two genetic approaches – transgenesis and paratransgenesis – is the most effective? and 2) can the two approaches be combined to enhance the effectiveness of the intervention? We use transgenic mosquitoes engineered to express effector genes in the midgut and/or salivary glands and *Serratia* bacteria engineered to express the same effector genes. We measured the ability of these two strategies, individually and in combination, to inhibit malaria parasite transmission.

## Results

### Generation of *An. stephensi* mosquitoes expressing anti-malaria effectors

To constitutively and robustly express anti-malaria effector proteins in the midgut and salivary glands of *An. stephensi* mosquitoes, we used the QF-QUAS binary expression system previously adapted for expression in *An. gambiae*.^15, 16^ We constructed two “driver” mosquito lines that express the QF transcription factor, one driven by the constitutive salivary gland-specific (AAPP) promoter^17^ and the other driven by the constitutive midgut-specific peritrophin 1 (Aper1) promoter^18, 19^ (Figure 1A). We also constructed a third “effector” mosquito line that encodes two parasite inhibiting factors (MP2 and scorpine) downstream of the QUAS promoter and driven by the QF transcription factor (Figure 1A). Crossing this effector line with either, or both, driver lines, leads to the salivary gland and/or midgut expression of parasite-inhibiting factors. The MP2 (midgut peptide 2) dodecapeptide, identified from a phage display screen, binds tightly to the mosquito midgut epithelium and inhibits *P. falciparum* invasion with high efficiency^20^, whereas the scorpion (*Pandinus imperator*) peptide scorpine lyses malaria parasites without affecting mosquito fitness^21, 22^. Each of the three constructs also expresses YFP (yellow eyes, salivary gland QF driver), dsRed (red eyes, midgut QF driver), or CFP (blue eyes, QUAS effector) fluorescent selection markers (Figure 1A).

**Figure 1.**
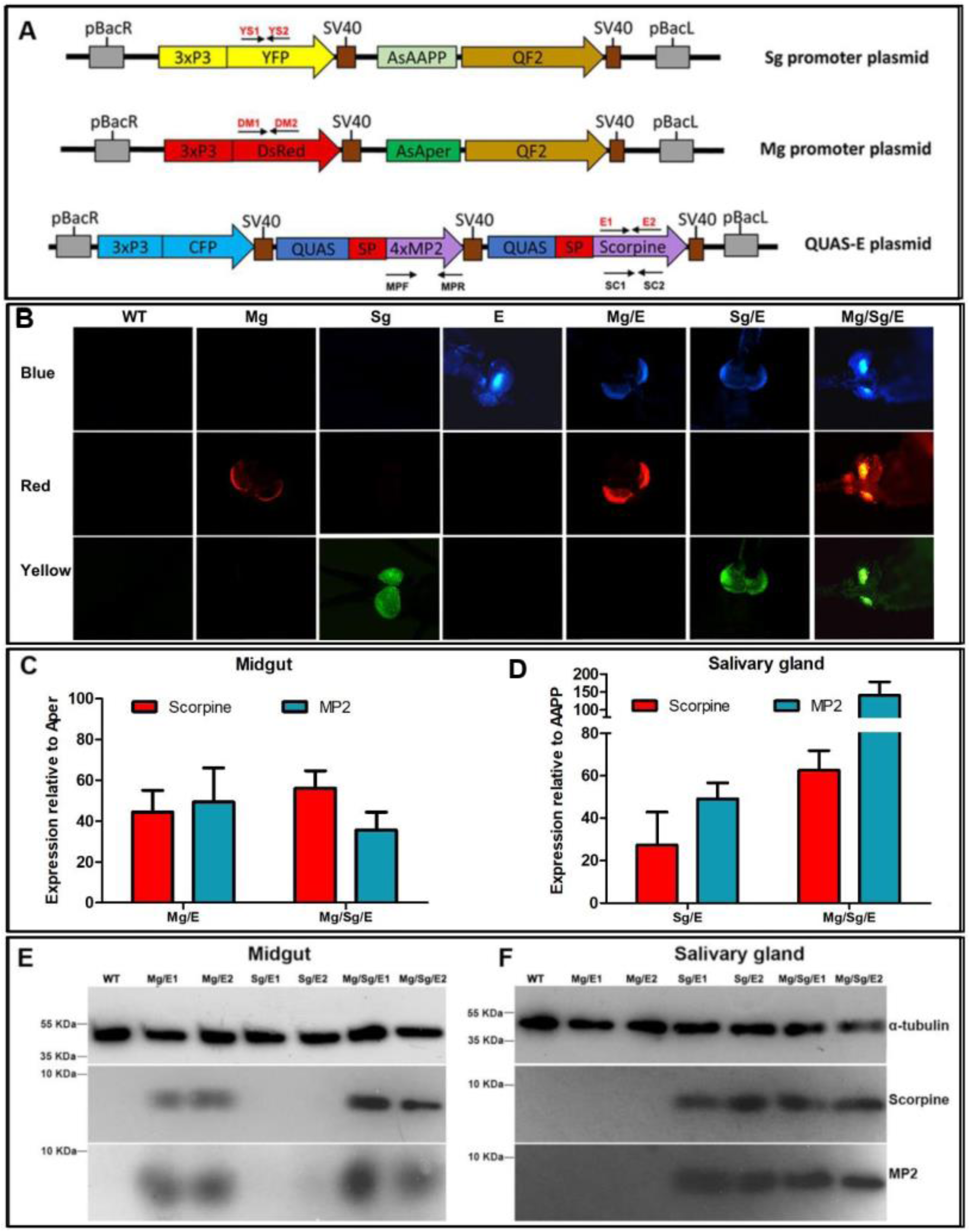
Tissue-specific expression of effector genes in *An. stephensi* transgenic mosquitoes. [**A]:** Diagram of the salivary gland (Sg) and midgut (Mg) driver constructs expressing the QF2 transcription factor and of the effector (E) constructs expressing the MP2 and scorpine effector proteins under control of the QUAS promoter. Each construct also includes sequences encoding a yellow (YFP), red (DsRed) or blue (CFP) fluorescent protein under the control of the 3xP3 eye promoter. pBac: piggyBac inverted terminal repeats; SV40: transcription terminator sequence; SP: *An. stephensi* carboxypeptidase signal peptide. Primers used for validation of insertion into mosquito lines (Figure S1 and Table S7) are indicated in red font. Primers used for qRT-PCR are indicated in black font (Table S7). [**B]** Detection of fluorescent eye markers in wild type (WT) and transgenic mosquitoes carrying different combinations of midgut driver (Mg), salivary gland driver (Sg) and effector (E) sequences. Tissue-specific expression of MP2 and scorpine mRNA in transgenic mosquitoes quantified by qRT-PCR in the midgut relative to the endogenous Aper mRNA **[C]**, and in the salivary glands relative to the endogenous AAPP mRNA **[D]**. Mosquito rpS7 was used as a reference. Data pooled from three independent biological replicates. Statistical analysis was determined by the Student′s t test. [**E]** and **[F]** Immunoblotting showing MP2 (6.17 kDa) and scorpine peptide (8.46 kDa) protein expression in midgut and salivary gland lysates from wild type and transgenic lines. α-tubulin was used as a loading control. E1 and E2 refer to independent mosquito transgenic lines. Antibodies used are shown to the right of [F].

Two midgut driver lines (Mg1 and Mg2), two salivary gland driver lines (Sg1 and Sg2) and two effector lines (E1 and E2) were obtained. Transgenic mosquitoes were screened by fluorescence microscopy (Figure 1B), and plasmid insertion was verified by PCR (Figure S1). The position of genome integration was determined for each parental line by splinkerette PCR^23^ and sequencing (Table. S1). All the parental lines, except for Sg2, have insertions in intergenic regions. Two of the three Sg2 insertions are in intergenic regions, and one in the open reading frame of the gamma-glutamyltranspeptidase gene (ASTE010947). Each transgenic line was propagated for over 10 generations, discarding at each generation mosquitoes not displaying the correct combination of fluorescent eyes, and this resulted in homozygous lines (Table S2).

### Quantification of effector mRNA and protein expression

Using reverse transcription quantitative polymerase chain reaction (RT-qPCR) we compared abundance of the endogenous mosquito Aper and AAPP transcripts with the abundance of effector transcripts originating from the same promoters but driven by the Q-system. Transcripts derived from the Q-system were substantially higher. In the midgut, the scorpine transcript was between 44- (*P*<0.01) and 56-fold (*P*<0.01) higher than that of the endogenous Aper mRNA and the MP2 transcript abundance was between 49- (*P*<0.001) and 36-fold (*P*<0.01) higher, depending on the transgenic line (Figure 1C; Table S3). In the salivary glands, scorpine transcript abundance varied between 27- (*P*<0.05) and 63-fold (*P*<0.01) higher and MP2 transcript between 49- (*P*<0.001) and 140-fold (*P*<0.01) higher than that of the endogenous AAPP mRNA, depending on the transgenic line (Figure 1D; Table S4). Moreover, in the absence of a driver, transgene expression in ‘E’ effector mosquitoes (see Figure 1A) was undetectable (Tables S3 and S4). These results attest to the high effectiveness of the Q-system in enhancing tissue-specific transgene expression. Western blot analysis using anti-MP2 and anti-scorpine antibodies confirmed the tissue specific expression of the MP2 (6.17 kDa) and scorpine (8.46 kDa) proteins (Figure 1E and F).

### Mosquito fitness is not affected by effector gene expression

To determine whether DNA integration or anti-malaria effector expression affects mosquito fitness, we analyzed the survival of WT, parental transgenic (Mg, Sg, and E) and transgene-expressing mosquitoes. No significant longevity differences were detected for any female (Figure S2A) or male (Figure S2B) transgenic mosquitoes, compared to WT. Next, we determined the fecundity (number of laid eggs) and fertility (percentage of hatched eggs) of WT, parental, and anti-malaria transgenic lines. Mosquitoes from all parental and anti-malaria-expressing transgenic lines showed no difference in fecundity when compared to WT mosquitoes (Figure S2C). As for fertility, no significant differences were detected for the Mg and Sg/E lines when compared to WT, while only marginal differences were detected for the Sg, E, Mg/E and Mg/Sg/E lines (2.0%, 3.1%, 2.0% and 2.0% reduction, respectively) (Figure S2D). To determine if transgenesis or anti-malaria gene expression in the midgut and/or in the salivary glands affects blood feeding, we quantified the proportion of mosquitoes that take a blood meal (feeding rate) and the amount of blood ingested per mosquito. We found no significant differences (Figure S2E and S2F), suggesting that neither transgenesis nor anti-malaria gene expression affects blood ingestion.

In summary, our data show that transgenesis and anti-malaria gene expression in the midgut and/or salivary glands does not impair mosquito survival, fecundity, fertility (only minor differences), and blood feeding under laboratory conditions.

### Effector-expressing *Serratia* populate the mosquito reproductive organs and are transmitted vertically and horizontally

Previously we reported that fluorescently labelled *Serratia* AS1 can spread through mosquito populations and that the bacteria are inherited through multiple mosquito generations^14^. Here we tested whether a *Serratia* AS1 strain (termed *Serratia AS1*-poly - for poly-effector) that produces and secretes the same effector proteins (MP2 & scorpine, among others) as those produced by the transgenic mosquitoes can also populate mosquitoes and be transmitted from one mosquito generation to another. We fed WT and Mg/Sg/E transgenic female mosquitoes with *Serratia AS1*-poly bacteria and quantified their capacity to colonize different mosquito organs and to be transmitted along consecutive mosquito generations. We found that *Serratia AS1*-poly equally populate WT and transgenic mosquito midguts, ovaries and accessory glands and are transmitted for at least three generations (Figure 2A-D). Moreover, we found that WT and transgenic male mosquitoes colonized with *Serratia AS1*-poly transferred the bacteria horizontally (sexually) to virgin WT and transgenic female mosquitoes (Figure 2E). Horizontal transfer did not take place when male mosquitoes were placed with mated females, showing that transfer occurs during copulation (Table S5; female mosquitoes mate only once in their lifetimes). These results suggest that recombinant *Serratia* AS1 can effectively populate transgenic mosquitoes and be transmitted through multiple generations.

**Figure 2.**
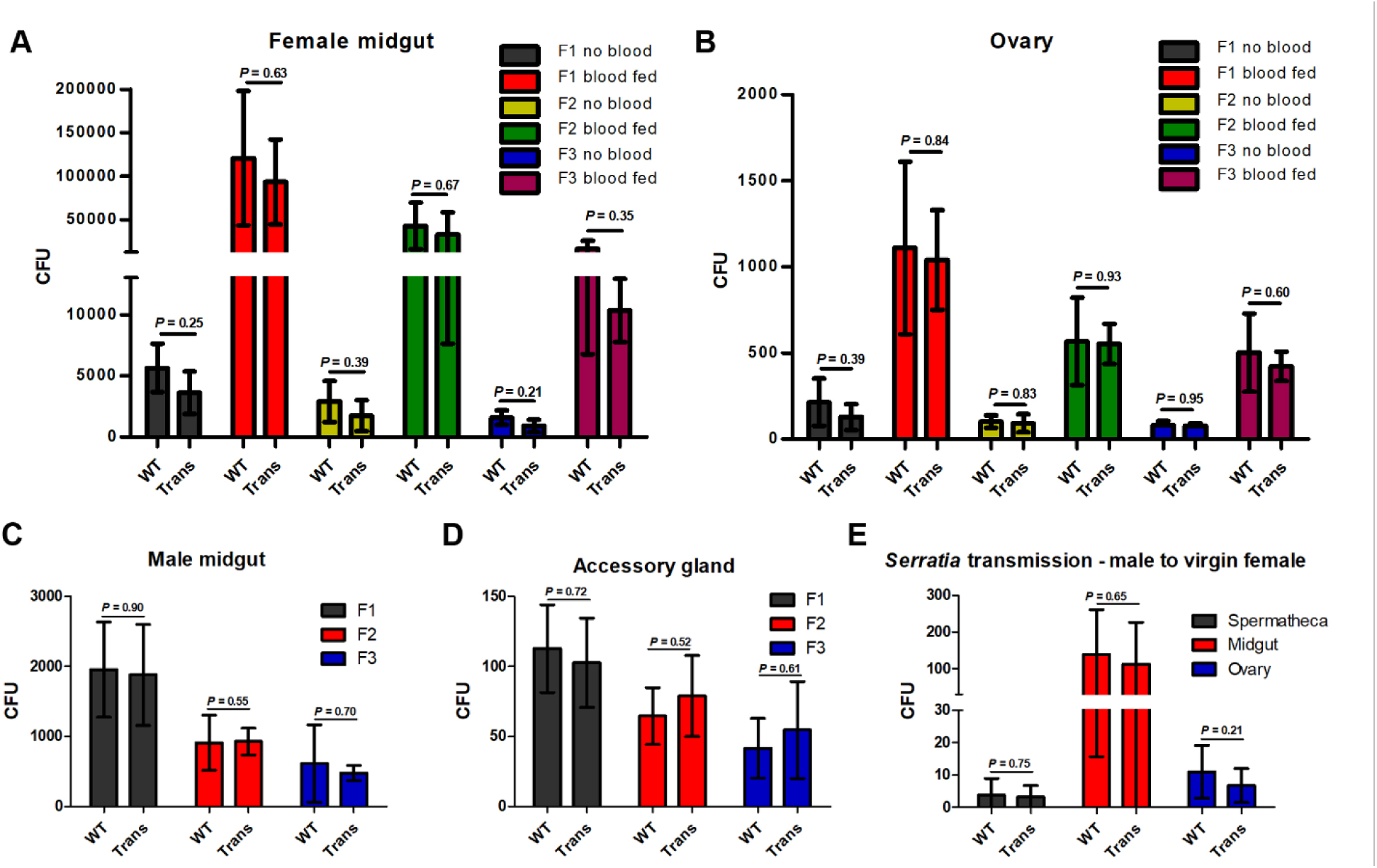
*Serratia AS1*-poly-effector bacteria persist through multiple mosquito generations. A total of 100 WT or transgenic (Trans) virgin females that had been fed with *AS1*-poly-effector bacteria were placed in a cage with 100 WT or transgenic virgin males (not fed with bacteria) and allowed to mate. Mosquitoes were then fed blood and allowed to lay eggs. These eggs were allowed to hatch and reared to adults following standard protocol (F1). The F1 mosquitoes were propagated through two additional generations (F2 & F3) without providing additional genetically modified bacteria. At each generation, 10 mosquitoes were dissected, and bacterial load was determined by plating serial dilutions of tissue homogenates on apramycin and ampicillin agar plates and counting colonies. **[A]** Colony-forming units (CFUs) per female midgut fed or not on blood. **[B]** CFUs per female ovary fed or not on blood. **[C]** CFUs per male midgut. **[D]** CFUs per male accessory gland. Data pooled from 3 independent experiments. **[E]** *Serratia* horizontal (sexual) transmission. Newly emerged virgin male adult mosquitoes were fed on 5% sugar solution containing 10^7^/ml *AS1*-poly-effector bacteria/ml and then allowed to mate with virgin females. Three days later, 10 females were assayed for the presence of *Serratia AS1* by plating spermatheca, midgut and ovary homogenates on apramycin and ampicillin agar plates and counting colonies. Trans: Mg/Sg/E transgenic mosquitoes. Error bars indicate standard deviation of the mean. Data pooled from 3 independent biological experiments. Statistical analysis was determined by the Student′s t test. *P*>0.05: not significant.

A concern is the possibility of *Serratia* carried by the mosquito being incorporated into the mosquito salivary glands and being delivered with the bite of a mammalian host. To address this concern, mosquitoes previously fed with fluorescently labeled *Serratia* were allowed to feed on blood using a membrane feeder. The remaining blood in the feeder was collected, grown overnight in LB medium and plated. No bacteria were detected (Figure S3), even though the presence of even a single bacterium in the blood from the feeder would have been easily detected.

### Transgenic and paratransgenic expression of MP2 and scorpine inhibit *Plasmodium* development in the mosquito

Wild type or transgenic mosquitoes, carrying or not wild type or recombinant bacteria, were fed with the same *P. falciparum* infectious blood. Infections were followed by measuring the formation of midgut oocysts (Figure 3A) and salivary gland sporozoite numbers (Figure 3B). Expression of effector molecules in the midgut or in the salivary glands of transgenic mosquitoes significantly reduced parasite burden, whereas concomitant effector expression in both organs reduced burden to the greatest extent (81% and 85% inhibition of mean oocyst and sporozoite numbers, respectively). Effector-expressing recombinant bacteria also significantly reduced parasite burden in WT mosquitoes (70% and 65% inhibition of oocyst and sporozoite numbers, respectively). Importantly, combining mosquito transgenesis with paratransgenesis led to the strongest inhibition of parasite development. Oocyst prevalence was reduced from 98% to 49% for transgenic-only mosquitoes and to 48% when transgenesis and paratransgenesis were combined. Sporozoite prevalence was reduced from 97% to 42% for transgenic-only mosquitoes and to 24% when transgenesis and paratransgenesis were combined. These results suggest that the ability of [transgenic + paratransgenic] mosquitoes to transmit the parasite may be strongly impaired, a hypothesis that was tested next.

**Figure 3.**
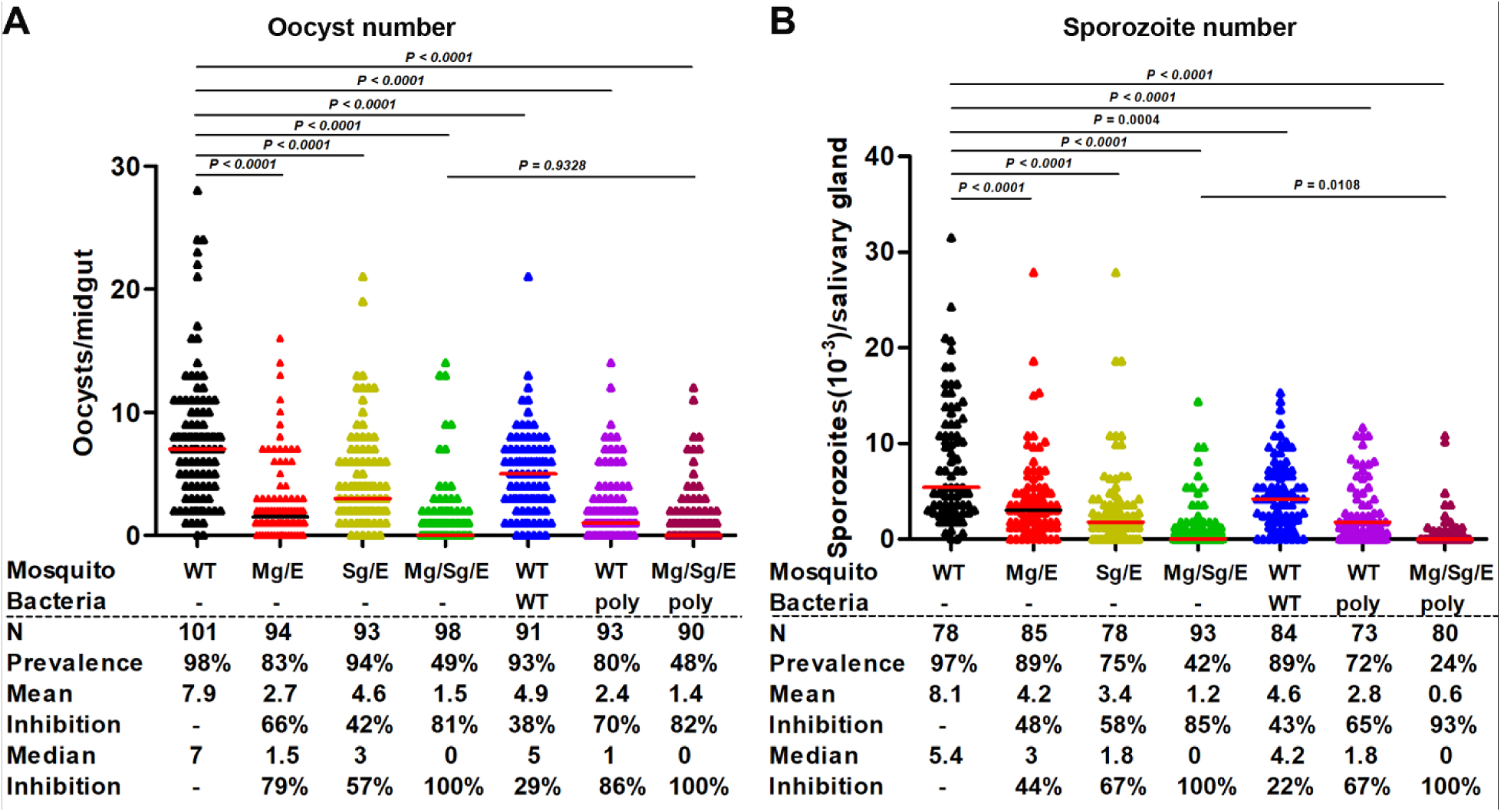
Transgenesis and paratransgenesis strongly impair *Plasmodium* development. Two-day-old *An. stephensi* mosquitoes were fed (or not) overnight with wild type or recombinant *Serratia* AS1 (poly) bacteria, as indicated. After 48 h, all mosquito groups were fed on the same *P. falciparum* gametocyte culture and midgut oocyst number was determined on day 7 **[A]** and salivary gland sporozoite number was determined on day 14 **[B]** post-feeding. Horizontal red lines represent median oocyst or sporozoite number. Data pooled from three independent biological experiments. Statistical analysis by Mann–Whitney U test. *P*> 0.05: not significant; ‘poly’: *Serratia AS1*-poly bacteria; N: number of mosquitoes assayed; Prevalence: proportion of mosquitoes carrying one or more parasite.

### Malaria transmission is maximally impaired by combining transgenesis and paratransgenesis

To investigate the ability of mosquitoes to transmit the parasite from an infected to a naïve animal, we challenged naïve mice with the bite of mosquitoes that had ingested the same infectious blood meal. Four mosquito groups were investigated: 1) wild type mosquitoes (WT), 2) WT mosquitoes carrying *Serratia* AS1-poly (paratransgenic), 3) transgenic mosquitoes that express effectors in the midgut and salivary glands (transgenic) and 4) transgenic mosquitoes carrying *Serratia* AS1-poly (paratransgenic + transgenic) (Figure 4A). All four mosquito groups ingested the same number of parasites, as they were fed on the same *P. berghei*-infected mouse. At 21-23 days post-feeding, after the mosquito salivary glands were populated by sporozoites, either three (Figure 4B) or five (Figure 4D) mosquitoes were randomly selected and allowed to bite naïve mice. For each experiment, salivary gland sporozoite numbers were determined (Figure 4C and 4E).

**Figure 4.**
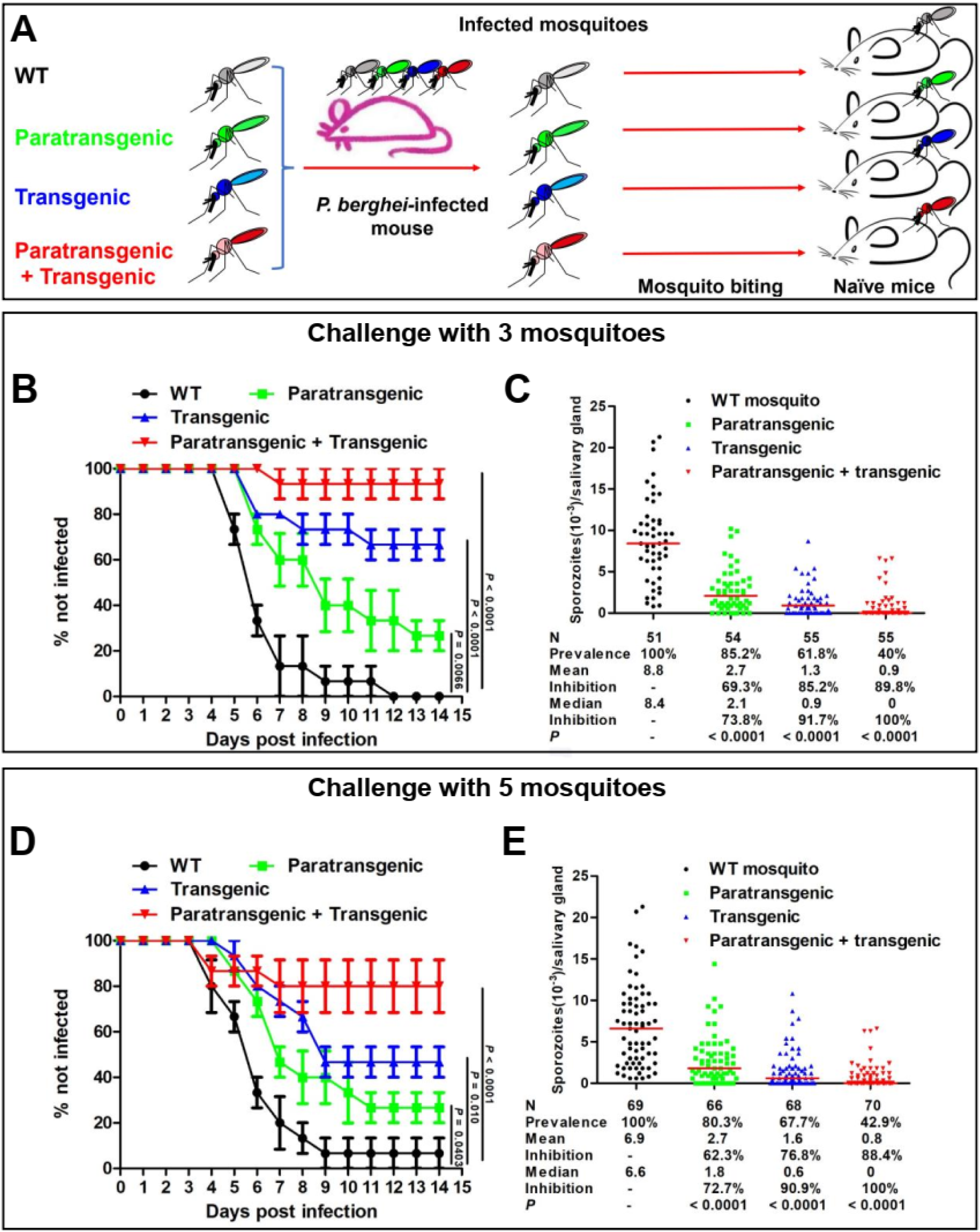
Transgenesis and paratransgenesis inhibit *P. berghei* transmission by mosquitoes from infected to naïve mice. **[A]** Experimental design. Wild type (WT), paratransgenic, transgenic and [paratransgenic + transgenic] mosquitoes were fed on the same *P. berghei*-infected mouse, assuring that all mosquitoes ingested the same number of parasites. After 21∼23 days, when sporozoites had reached the salivary glands **[C, E]**, three **[B]** or five **[D]** mosquitoes were randomly selected and allowed to bite naïve mice. The parasitemia of these mice was followed for 14 days. Data pooled from three independent experiments, each using five mice per challenged group for a total of 15 mice. Transgenic mosquitoes express effectors in both midgut and salivary glands. Data pooled from three independent biological experiments. Statistical analysis (B and D) was determined by the Log-rank (Mantel-Cox) test; Statistical analysis (C and E) was determined by the Mann–Whitney U test.

When mice were challenged with the bite of three WT mosquitoes (three independent experiments with five mice each), 100% became infected (half-infection time = 5.5 ± 0.5 d) (Figure 4B) and their salivary glands had a median 8,400 sporozoites (Figure 4C). With mosquitoes carrying *Serratia* AS1-poly (paratransgenic), 26.7% of the mice were protected (half-infection time 7.1 ± 0.7 d), and their salivary glands had a median of 2,100 sporozoites (74% lower than WT mosquitoes). With transgenic mosquitoes, 67% of the mice were protected, and their salivary glands had a median of 900 sporozoites (92% lower than WT mosquitoes). With [paratransgenic + transgenic] mosquitoes, 93% mice were protected, and their salivary glands had a median of zero sporozoites (100% lower than WT mosquitoes).

When mice were challenged with the bite of five WT mosquitoes (Figure 4D and 4E), only one mouse out of 15 (6.7%) did not get infected (half-infection time = 5.6 ± 0.7 d). Paratransgenesis, protected 26.7% mice (half-infection time = 8.3 ± 1.0), transgenesis protected 47% mice (half-infection time = 10.1 ± 1.0 d) and [paratransgenesis + transgenesis] protected 80% mice. The salivary gland sporozoite number (Figure 4E) was similar to that observed for experiments with three mosquito bites (Figure 4C).

In summary, our data shows that transgenic and paratransgenic expression of MP2 and scorpine are both effective in impairing transmission, and that the combination of the two complementary strategies is considerably more effective. An even higher protection is expected from the bite of one infected mosquito, which is the most likely scenario in the field.

## Discussion

In this study, we report the development of transgenic mosquitoes and paratransgenic bacteria expressing anti-plasmodial effector molecules and the effect of these two technologies, individually or in combination, on the transmission of *Plasmodium* parasites. The Q-binary expression system was used to express effector genes in the mosquito midgut and salivary glands. Notably, effector mRNA abundance was about 50-times higher than that of the endogenous genes, consistent with the high effectiveness of the Q-system in *Drosophila*.^15^ Some QF toxicity was reported when the Q-system was first used in *Drosophila*.^15, 16^ Of note, expression of neither the QF transcription factor nor the anti-malaria effectors affected mosquito longevity, blood meal uptake or offspring production under laboratory conditions. Additional experiments are required to test the fitness of these transgenic mosquitoes under field conditions.

We selected two potent effectors molecules, MP2 and scorpine, to block the development of *Plasmodium* in the mosquito. MP2 is a 12-amino-acid peptide that likely targets a midgut receptor for ookinete traversal,^20^ and scorpine is an antimicrobial toxin hybrid between a cecropin and a defensin, that lyses *Plasmodium* ookinetes.^21^ Scorpine expressed by the entomopathogenic fungus *Metarhizium* in the mosquito hemocoel strongly inhibits (∼90%) salivary gland sporozoite numbers.^24^ Transgenic mosquitoes expressing this effector in the salivary glands were also highly effective in reducing sporozoite numbers (this work). Furthermore, expression of both effector genes in the midgut and the salivary glands led to a much stronger decrease of salivary gland sporozoite numbers than the expression of the effectors in either of these organs alone. A number of effectors have already been individually tested in paratransgenesis experiments.^14^ Going forward, the combination of different effectors and the use of mosquitoes and bacteria expressing different effector sets should be explored, to achieve maximum blocking activity.

That expression of anti-malaria effectors in the salivary glands inhibited oocyst development in the midgut is most likely explained by the fact that mosquitoes ingest saliva with the blood meal, in this way incorporating effector proteins into the blood bolus.^25^ Effector molecules secreted in the saliva could conceivable be injected into the dermis of the host during blood feeding. Scorpine has been shown to be non-toxic to insect cells,^26^ whereas MP2 toxicity has not been determined. However, their toxicity has not been tested in the presence of mosquito saliva. Therefore, further studies are needed to determine if the delivery of these molecules by the mosquito bite could induce physiological responses.

Experiments seeking evidence for possible bacteria transmission with a mosquito bite yielded negative results (Figure S3), suggesting that mosquitoes cannot inoculate the bacteria while feeding on a host. It was previously shown that secretion of effector proteins by recombinant *Pantoea*,^6^ *Serratia*^14^ or *Asaia*^12^ bacteria into the midgut inhibits *Plasmodium* development and that *Serratia* AS1 is transmitted from one mosquito generation to the next.^14^ What was not known is whether engineering *Serratia* to produce and secrete large amounts of proteins would affect their fitness and ability to be transmitted. Our experiments showed that the engineered *Serratia* were efficiently transmitted from one mosquito generation to the next, a result that bodes well for the implementation of the paratransgenesis strategy in the field.

This project was based on two basic premises: (i) transgenesis and paratransgenesis are not mutually exclusive and (ii) both strategies result in impairment of parasite development in the mosquito. As such, our experiments addressed the question of whether a combination of the two strategies would result in enhanced transmission-blocking effectiveness. The combination of transgenesis and paratransgenesis greatly reduced parasite development in the mosquito and most importantly, it resulted in a high-level reduction of transmission from an infected to a naïve mouse, compared to individual interventions. When mice were bitten by three [transgenic + paratransgenic] mosquitoes, 93% of the mice were protected from infection as compared with zero protection when mice were bitten by WT mosquitoes that acquired the parasites from the same infected mouse. In the field, where the density of infected mosquitoes is low even in high-transmission areas, it is unlikely that people will be consecutively bitten by more than one infected mosquito, and protection from transmission is expected to be very high. For translating these findings to the field, the testing of different combinations of effectors, both for transgenesis and paratransgenesis, may further improve the effectiveness of the approach.

Whereas both transgenesis and paratransgenesis have been shown to be highly effective in a lab setting, the challenge is to implement this new containment strategies in the field. In addition to address regulatory and ethical issues connected with the release of recombinant organisms in nature, a major technical issue to be solved is how to introduce the blocking transgenes into mosquito populations in the field. In this respect, CRISPR/Cas9 technology has afforded the development of promising gene drive systems^27, 28, 29^ focused on population suppression or population modification strategies. Population reduction leaves an empty biological niche that upon cessation of reduction pressure, will result in recolonization by the same or other mosquito species. In contrast, population modification results in a more stable state, with a biological niche occupied by mosquitoes that are poor transmitters. Similarly, efficient spread of recombinant bacteria into mosquito populations has been demonstrated in a laboratory setting (,^14^ this work), indicating a promising path toward the field implementation of the most efficient [transgenesis + paratransgenesis] strategy. The recent finding that a naturally occurring and non-modified *Serratia* can spread through mosquito populations while strongly suppressing *Plasmodium* development,^30^ significantly increases the feasibility of moving paratransgenesis into the field, as it bypasses concerns relating to the release of genetically modified organisms in nature. In the field, we envision the use of attractive sugar feeding stations for *Serratia* introduction into mosquito populations.^31^ Female mosquitoes that acquire the bacteria will seed the breeding sites when they lay eggs.^14^ Notably, transgenesis and paratransgenesis are not envisioned to be implemented by themselves. Both are compatible with current vector and malaria control measures such as insecticide-based mosquito control, mass drug administration, and vaccines, and their added implementation promises to substantially enhance the effectiveness of intervention of disease transmission.

In summary, we show that the Q-binary system to express anti-*Plasmodium* effectors in the mosquito is highly efficient. We also show that in addition to inhibiting parasite development, recombinant *Serratia* AS1 is horizontally and vertically transmitted across multiple mosquito generations, which is a bacteria counterpart of gene drive. A major conclusion of this work is that the combination of transgenesis with paratransgenesis provides maximum parasite blocking activity and has high potential for fighting malaria.

## Material and Methods

### Animal Handling and Ethics Protocol

#### Ethics statement

This study was carried out in accordance with the guidelines of the Johns Hopkins University Animal Care and Use Committee (AUCC) under protocol number: M018H18.

#### Mosquitoes rearing and parasite culture

*Anopheles stephensi* Nijmegen strain^32^ and *An. stephensi* transgenic lines were reared as previously described^33^. For fitness evaluation, the mosquitoes were fed on Swiss Webster mice.

Female *An. stephensi* were infected with *P. falciparum* gametocyte cultures via membrane feeding. *P. falciparum* NF54 gametocytes were produced according to Tripathi et al.,^34^. Briefly, the parasites were maintained in O+ human erythrocytes using RPMI 1640 medium supplemented with 25 mM HEPES, 50 mg/L hypoxanthine, 25 mM NaHCO3, and 10% (v/v) heat-inactivated type O + human serum (Interstate Blood Bank, Inc.) at 37 °C and with a gas mixture of 5% O_2_, 5% CO_2_, and balanced N_2_. For feeding, 14–17-day-old mature gametocytes were pelleted by centrifugation (5 min, 2,500 g), resuspended with O + human RBC to 0.15%-0.2% gametocytemia and diluted to 40% hematocrit with human serum. All manipulations were done maintaining the cultures, tubes, and feeders at 37 °C.

#### Plasmid constructs

The pXL-BACIIECFP-15XQUAS-TATA-MP2-SV40-15XQUAS-TATA-scorpine-SV40 containing the MP2 and Scorpine expression cassette and the ECFP gene under the eye-specific promoter 3xP3 was used to generate the parental QUAS-[MP2+scorpine] effector lines (Table S6). The coding DNA for MP2-SV40-15XQUAS-TATA-Scorpine was synthetized by GeneScript (Figure S4). The sequence was amplified using primers MP2-ScopineF and MP2-ScopineR (Table S7), and In-Fusion-cloned into plasmid pXL-BACIIECFP-15XQUAS-TATA-SV40^15^ previously linearized with XhoI.

The pXL-BACII-DsRed-AsAper-QF2-hsp70 containing the QF2 transcription factor under the control of the midgut specific AsAper promoter and the DsRed marker driven by the eye-specific promoter 3xP3 was used to generate the parental Mg-QF driver line. The AsAper promoter (1.5 kb) (Figure S4) was PCR amplified from *An. stephensi* gDNA with primers MgPF and MgPR (Table S7). The PCR product was In-Fusion-cloned into plasmid pXL-BACII-DsRed-QF2-hsp70^15^ previously linearized with XhoI.

The pXL-BACII-YFP-AsAAPP-QF2-hsp70 containing the QF2 transcription factor under the control of the midgut specific AsAAP promoter and the YFP marker driven by the eye-specific promoter 3xP3, was used to generate the parental Sg-QF driver lines. The YFP coding sequence was amplified using primers YFPF and YFPR (Table. S7) (Figure S4). The PCR product was In-Fusion-cloned into plasmid pXL-BACII-DsRed-QF2-hsp70 previously digested with ApaI and NotI to produce plasmid pXL-BACII-YFP-QF2-hsp70. The AsAAPP promoter consisting of a 1.7 kb upstream of the start codon^19^ was PCR-amplified from *An. stephensi* gDNA using primers SgPF and SgPR (Table. S7) (Fig. S4). The PCR product was In-Fusion-cloned into plasmid pXL-BACII-YFP-QF2-hsp70 previously linearized with XhoI.

#### Generation of transgenic mosquitoes

The plasmid constructs were microinjected into *An. stephensi* embryos as described.^35^ Briefly, transformation plasmids were purified using the EndoFree Maxi Prep Kit (Qiagen) and resuspended in injection buffer (0.1 mM NaHPO4 pH 6.8 and 5 mM KCl) at a concentration of 250 ng/µl for the transformation plasmid and 200 ng/µl for the helper plasmid encoding the transposase. The plasmid mix was injected into *An. stephensi* embryos using a FemtoJet Microinjector (Eppendorf). Third instar larvae of G_0_ survivors were screened for transient expression of the 3xP3-dsRed marker (red eyes), 3xP3-YFP marker (yellow eyes), 3xP3-CFP marker (blue eyes). Adults obtained from the fluorescent marker screening were crossed to WT mosquitoes to generate independent transgenic lines. The data for these injections are summarized in Table S8.

For each of the parental transgenic lines, splinkerette PCR^23^ and PCR sequencing were used to determine the transgene insertion site into the *An. stephensi* genome. (Two rounds of amplifications were conducted with 1X Phusion High-Fidelity PCR Master Mix with HF Buffer (Thermo Fisher Scientific). The primers used are shown in Table S7. The amplified PCR products were resolved in a 1.5% agarose gel stained with ethidium bromide, and the amplified DNA bands from the 5’ and 3’ ends were individually excised and purified with QIAquick^®^ Gel Extraction Kit (QIAGEN). Purified PCR products were cloned into pJET1.2/blunt plasmid (Thermo Fisher Scientific) and transformed into NEB 5-alpha Competent *Escherichia coli* (High Efficiency, Thermo Fisher Scientific). Plasmids were isolated from individual colonies and sequenced with the universal primers pJET12F and pJET12R (Eurofins). The sequences were aligned to the *An. stephensi* genome using VectorBase and NCBI BLAST to identify the location of transgene insertion sites (Figure S1).

To obtain homozygous lines, each transgenic line was propagated for more than 10 generations, discarding at each generation mosquito larvae not displaying the expected fluorescent eyes. To verify homozygosity of the transgenic lines, 10 females of each line were mated with 10 WT male mosquitoes, fed blood, eggs were collected and reared to larvae. The larvae were individually inspected for expression of the fluorescent protein marker(s). Absence of the expected fluorescence would indicate that the parent female was heterozygous for this dominant marker.

To induce midgut- or salivary gland-specific expression of MP2 and scorpine, QF driver lines were crossed to QUAS-[MP2+scorpine] effector lines. The offspring of each cross was selected by the specific combination of eye fluorescence reporters (Figure 1B).

#### Quantitative reverse transcription polymerase chain reaction (qRT-PCR)

Tissue specific expression of MP2 and scorpine mRNAs in *An. stephensi* transgenic lines was evaluated by RT-PCR. Salivary glands and midguts were dissected from female mosquitoes in ice-cold 200 µl TRIzol® (Thermo Fisher Scientific). Total RNA was extracted according to TRIzol® manufacturer’s protocol, resuspended in RNAse free water, and treated with RQ1 RNase-Free DNase® (Promega; Madison, WI, USA). After RNA quantification using a DeNovix DS-11 spectrophotometer, first-strand cDNA was synthesized for each sample using Superscript III (Invitrogen) with random hexamers (Invitrogen) and 500 ng of total RNA per sample. cDNA was treated with RNase H (New England Biolabs) for 10 min at 37 °C and stored at -70 °C until use. The cDNA was used as template in PCR reactions containing the Taq 2X Master Mix (New England Biolabs) and 5 μM of MP2- and scorpine-specific primers (Table S7). Amplification of S7 ribosomal mRNA was used as reference.^36^ PCR conditions were: 1 hot start at 95 °C for 30 sec; 35 cycles of denaturation at 95 °C for 30 sec, annealing at 56 °C for 30 sec, and elongation at 68 °C for 30 sec; followed by a final extension at 68 °C for 5 min; and 4 °C indefinitely.

#### Mice immunization

Scorpine epitope (CEKHCQTSGEKGYCHGT, the N-terminus was conjugated to KLH) and MP2 epitope (ACYIKTLHPPCS, the N-terminus was conjugated to KLH) were synthesized by Peptide 2.0 Inc. About 6-8-week-old C57BL/6 mice were immunized with 20 µg (50 µl) purified antigen in PBS using Addavax (Invivogen, San Diego, CA) as the adjuvant. A total of 50 µl adjuvant was mixed with 50 µl antigen, and the mixture was administered intramuscularly in both anterior tibialis muscles (50 µl per leg). Mice were immunized twice at two-week intervals. Serum was collected 14-21 days after administration of the last booster.^37^

#### Commercial antibodies

Rabbit anti-α-tubulin was purchased from Sigma (cat# SAB3501072) and goat anti-rabbit IgG HRP-conjugated and goat anti-mouse IgG HRP-conjugated were purchased from Cell Signaling (cat# 7076S).

#### Western blotting

MP2 and scorpine protein synthesis in midgut and salivary glands of the transgenic lines was evaluated by Western blot. Five midguts and ten salivary glands were dissected in PBS and placed in microtubes containing RIPA Buffer® (Thermo Fisher Scientific), 1% Halt™ Protease Inhibitor Cocktail (Thermo Fisher Scientific), and 0.1 mM PMSF (Sigma-Aldrich). Samples were homogenized and stored at -70 °C. An equivalent of 0.25 midgut and 5 salivary glands were resolved in a NuPAGE™ 10% Bis-Tris Protein Gel (Invitrogen) under reducing conditions and transferred to a PVDF membrane Invitrogen™ Power Blotter Select Transfer Stacks. After the transfer, the membrane was washed with TBST 1% (Sigma-Aldrich), incubated with blocking buffer (5% milk powder in TBST 1%) overnight at 4 °C, and probed with mouse anti-MP2 or anti-scorpine at a 1:1,000 dilution in TBST 1% overnight at 4 °C. The membrane was washed and incubated with an anti-mouse HRP-linked antibody (Cell Signaling) at a 1:10,000 dilution in TBST 1% for 2 h at room temperature. Detection was done with the SuperSignal™ West Dura Extended Duration Substrate Chemiluminescent Substrate (Thermo Fisher Scientific), and imaged using an Azure Imager c600^®^ (Azure Biosystems).

#### Mosquito survival, fecundity, and fertility

To measure mosquito survival, two-day-old adult male and female mosquitoes (n = 100) were separately placed in a cage with cotton pads soaked in 10% sucrose solution and kept in the insectary. Female mosquitoes were allowed to blood feed on an anesthetized mouse for 30 min and allowed to lay eggs. Mortality of female and male mosquitoes was monitored 3 times per week. The differences among the survival curves (three independent replicates) were analyzed with the Log-rank (Mantel-Cox) test, using the WT as controls.

To assess fecundity (number of laid eggs) and fertility (percentage of hatched eggs), two-day-old adult females were blood-fed on anesthetized mice for 30 min. Only fully engorged females were used for these experiments. Two days after blood-feeding, 20 females were individually placed in 50 ml tubes containing a small cup with filter paper soaked in 2 ml of distilled water as a oviposition substrate. After three days, the filter papers with eggs were removed, and the number of eggs per mosquito was counted using a dissecting microscope. After counting, the eggs were placed in paper cups with 50 ml of distilled water to allow hatching. fertility was determined as the number of larvae divided by the total number of eggs. Fecundity and fertility of the transgenic lines were compared to WT mosquitoes, and all the experiments were repeated for a total of three biological replicates.

#### Quantification of blood uptake

The amount of blood ingested by *An. stephensi* transgenic mosquitoes was determined by measuring the amount of protein-bound heme detected in the mosquito midgut after a blood meal.^38^ Transgenic and WT mosquitoes were fed with a 1:1 mixture of plasma and RBCs (Interstate Blood Bank Inc.) using membrane feeders. After feeding, the midguts of ten fully engorged females were dissected and homogenized individually in 1 ml of distilled water. Unfed mosquitoes were used as the negative control. Protein-bound heme (410 nm) was measured for each individual midgut with a Versa max microscope Reader and recorded with Softmax pro 5.3. Readings were compared among the groups using Student’s t test.

#### Bacteria administration to *An. stephensi* mosquitoes

After culturing at 28 °C overnight, bacteria were washed with sterile PBS and resuspended to a final concentration of 10^9^/ml. After a 3 h starvation, mosquitoes were fed overnight on 10^7^ CFU bacteria (*AS1*-poly, apramycin resistance) per ml of 5% sugar. Mosquitoes were surface-sterilized with cold 75% ethanol for 3 min and washed three times with sterile PBS. Midguts were dissected under sterile conditions at different time points before and after a blood meal and homogenized in sterile PBS. Bacterial number was determined by plating ten-fold serial dilutions of the homogenates on LB agar plates containing 50 µg/ml apramycin and ampicillin (bacteria from non-infected mosquitoes cannot grow on LB agar plates containing 50 µg/ml apramycin and ampicillin) and incubating at 28 °C for 24 h.

#### Effect of bacteria on mosquito infection by *Plasmodium falciparum*

*Serratia* bacteria were administered overnight to female *An. stephensi* with a cotton pad soaked with a 5% sucrose solution containing 10^7^ bacteria/ml or no bacteria, and 2 d later, allowed to feed on *P. falciparum* NF54 gametocyte-containing blood as described.^24^ Engorged mosquitoes were kept at 27 °C and 80% relative humidity. Midguts were dissected in 1× PBS at 7 d post-infection, stained with 0.1% mercurochrome and oocysts were counted. Salivary glands from mosquitoes were dissected at 14 d post-infection and individually homogenized on ice in 30 µl of PBS using a disposable pestle. The homogenate was centrifuged at 2,000 rpm for 10 min to pellet tissue debris. Then, 10 µl of the suspension was placed in a Neubauer counting chamber, waiting for at least 5 min to allow sporozoites to sediment to the bottom of the chamber. Sporozoites were counted using a Leica phase-contrast microscope. Parasite numbers among control and experimental groups were compared using the nonparametric Mann–Whitney test (GraphPad, Prism).

#### Effect of bacteria on mosquito infection by *Plasmodium berghei*

Bacteria were cultured overnight in LB medium and washed three times with sterile PBS. Two-day-old mosquitoes were fed overnight on a cotton pad soaked with a 5% sucrose solution containing or not 10^7^ bacteria/ml. Two days later, mosquitoes were fed on a *P. berghei*-infected mouse (1-2% of parasitemia and 1 exflagellation per 10 fields). Unfed mosquitoes were removed, and fully engorged mosquitoes were provided with 5% (wt/vol) sterile sucrose solution and maintained at 19 °C and 80% relative humidity. Midguts were dissected on day 12 after the blood meal, stained with 0.1% (wt/vol) mercurochrome for determining oocyst load. Salivary glands were dissected at 21 d post-infection for sporozoite determination. Transgenic and WT mosquitoes were simultaneously fed on the same *P. berghei*-infected mouse to assure that control and experimental mosquitoes ingested the same number of parasites.

#### *Serratia* vertical, venereal and transstadial transmission

To test vertical transmission, *AS1*-poly were introduced into two-day-old adult female mosquitoes by feeding them overnight on a cotton pad moistened with 5% sterile sucrose containing 10^7^ bacteria/ml. Two days later, mosquitoes were fed on a healthy mouse and were then allowed to lay eggs on a damp filter paper in individual oviposition tubes. Eggs were collected into a tube containing 300 μl sterile 1×PBS and homogenized. The bacterial load was determined by plating ten-fold serial dilutions of the egg homogenates on LB agar plates containing 50 µg/ml of apramycin and ampicillin and incubating the plates at 28 °C for 24 h for colony counting. Rearing of larvae to adults followed standard protocol. A total of 10 male and 10 female adults were sampled and examined by plating adult midgut homogenates on LB agar plates containing apramycin and ampicillin. To test the efficiency of *Serratia* transmission through multiple generations, the mosquitoes were reared without providing additional *Serratia* AS1 and maintained for three consecutive generations. At each generation, 10 female and male adults were sampled for examining the presence of *AS1*-poly effectors.

For male-to-female venereal transmission tests, *Serratia* were introduced into newly emerged virgin male mosquitoes by feeding them overnight on a cotton pad moistened with 5% sugar solution containing 10^7^ bacteria/ml. Twenty *Serratia*-carrying males were then allowed to mate with 20 three-day-old virgin females. Three days after mating, 10 females were sampled and examined for bacteria in the female midgut, ovary and spermatheca.

#### Transmission from infected to naïve mice

Transgenic and WT mosquitoes were simultaneously fed on the same *P. berghei*-infected mouse and unfed or partially fed mosquitoes were removed. Midguts from a small number of mosquitoes were dissected at 12 d post-feeding to determine the infection status by counting oocyst numbers. At ∼21-23 days post-feeding, three or five mosquitoes were randomly selected from the cage and allowed to feed on non-infected mice (challenge). Mosquitoes that did not take a blood meal were replaced until the final number of mosquitoes for each group (three or five) was reached. The salivary glands of most mosquitoes were dissected for counting sporozoites. A total of five mice were used per experiment and three biological replicates were conducted for a total of 15 mice per mosquito group. After mosquito challenge, mice were monitored daily for 14 d to determine blood-stage infection using Giemsa-stained blood smears.

## Acknowledgments

We thank the Insectary and Parasite Core Facilities of the Johns Hopkins Malaria Research Institute. This work was supported by a grant R01AI031478 from the National Institutes of Health, by the NIH Distinguished Scholars Program, and the Intramural Research Program of the Division of Intramural Research AI001250-01, National Institutes of Allergy and Infectious Diseases, National Institutes of Health and by the Bloomberg Philanthropies. Supply of human blood was supported by the National Institutes of Health grant RR00052. We thank Dr. Yuemei Dong from the Johns Hopkins Malaria Research Institute for providing the helper plasmid and Dr. Christopher Potter from Johns Hopkins School of Medicine for providing pXL-BACIIECFP-15XQUAS-TATA-PAI-SV40 and pXL-BACII-DsRed-AAPP-QF2-hsp70 plasmids.

## Supplementary figures

**Figure S1.**
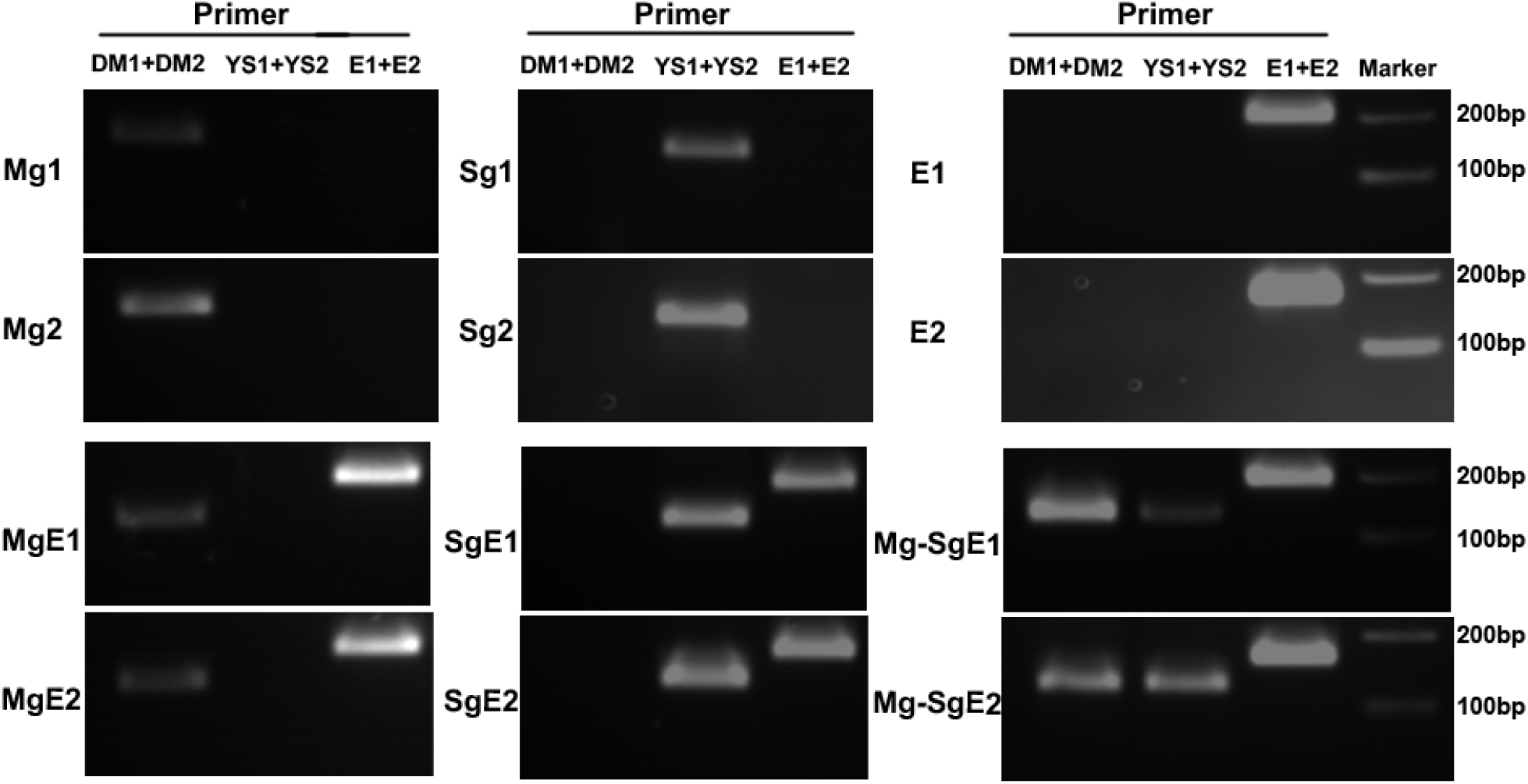
PCR validation of plasmid insertion site in mosquito lines. Primer pairs used for PCR reactions are indicated on top of each lane (sequences provided in Table S7; position of primers indicated in Figure 1A with red font). The DM1+DM2 primer pair was used to verify the MG QF2 driver plasmid insertion; the YS1+YS2 primer pair was used to verify the SG QF2 driver plasmid insertion; and the E1+E2 primer pair was used to verify the QUAS-MP2-QUAS-scorpine effector plasmid insertion. The transgenic lines are identified to the left of each panel.

**Figure S2.**
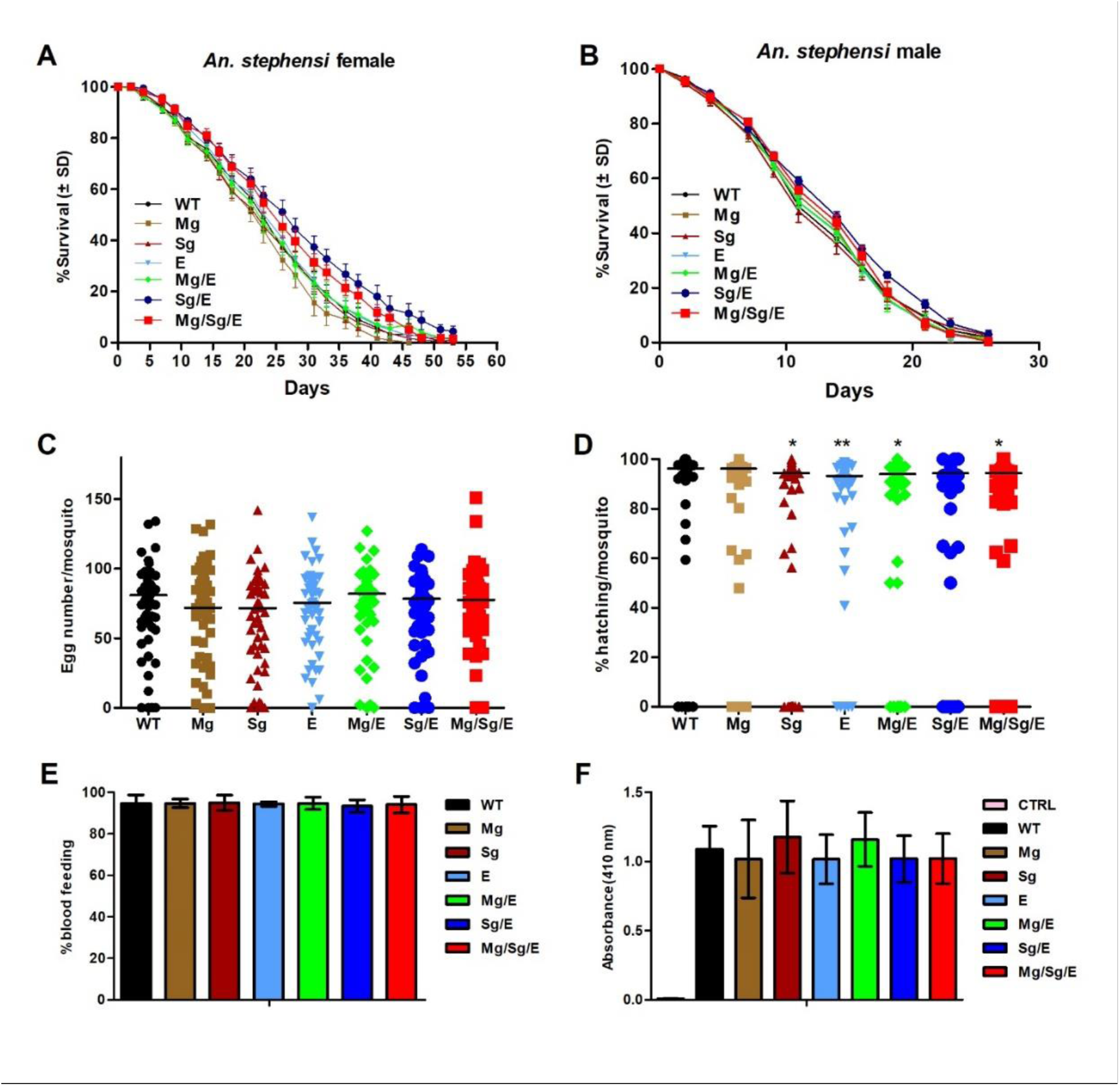
Fitness analysis of *An. stephensi* transgenic lines. **[A** and **B]:** Survival curves for WT and transgenic (see Figure 1A) females that received one blood meal on day two [A] and males [B], all maintained on sugar meal. No significant differences in survival rate were detected as calculated by Kaplan-Meier survival curves, and multiple comparisons by Log-rank test with Bonferroni correction for parental expressing lines. For [A] WT & Mg: *P* = 0.99; WT & Sg: *P* = 0.65; WT & E: *P* = 0.46; WT & Mg/E: *P* = 0.64; WT & Sg/E: *P* = 0.32; WT & Mg/Sg/E: *P* = 0.37 and for [B] WT & Mg: *P* = 0.66; WT & Sg: *P* = 0.53; WT & E: *P* = 0.99; WT & Mg/E: *P* = 0.97; WT & Sg/E: *P* = 0.075; WT & Mg/Sg/E: *P* = 0.34]. Combined from three biological replicates (N: 300 mosquitoes). **[C]:** Comparison of fecundity (number of laid eggs) between WT and transgenic mosquitoes. No significant differences were found using the Log-rank (Mantel-Cox) test. WT & Mg: *P* = 0.74; WT & Sg: *P* = 0.24; WT & E: *P* = 0.77; WT & Mg/E: *P* = 0.70; WT & Sg/E: *P* = 0.80; WT & Mg/Sg/E: *P* = 0.97 **[D]:** Comparison of fertility (proportion of laid eggs that hatched) between WT and transgenic lines. Statistical analysis used the Mann– Whitney U test. WT & Mg: *P* = 0.93; *WT & Sg: *P* = 0.021; **WT & E: *P* = 0.0089; *WT & Mg/E: *P* = 0.032; WT & Sg/E: *P* = 0.050; *WT & Mg/Sg/E: *P* = 0.032]. [C and D]: data combined from three biological replicates (N: 60 mosquitoes); horizontal lines are median values. **[E]** The percentage of mosquitoes that take a blood meal is not affected. Two-day-old female mosquitoes were allowed to feed on mice, and the percentage of mosquitoes that fed was determined after 30 min feeding. WT & Mg: *P* = 1.00; WT & Sg: *P* = 0.92; WT & E: *P* = 0.90; WT & Mg/E: *P* = 1.00; WT & Sg/E: *P* = 0.68; WT & Mg/Sg/E: *P* = 0.85. **[F]** The amount blood uptake is not affected. Quantification of protein-bound heme at 410 nm from midguts of WT and transgenic mosquitoes before (CTRL) and after a blood meal. WT & Mg: *P* = 0.73; WT & Sg: *P* = 0.65; WT & E: *P* = 0.63; WT & Mg/E: *P* = 0.66; WT & Sg/E: *P* = 0.64; WT & Mg/Sg/E: *P* = 0.76. [E and F]: error bars represent SD of the mean; data pooled from three independent experiments; o significant differences were found using the Student′s t test.

**Figure S3.**
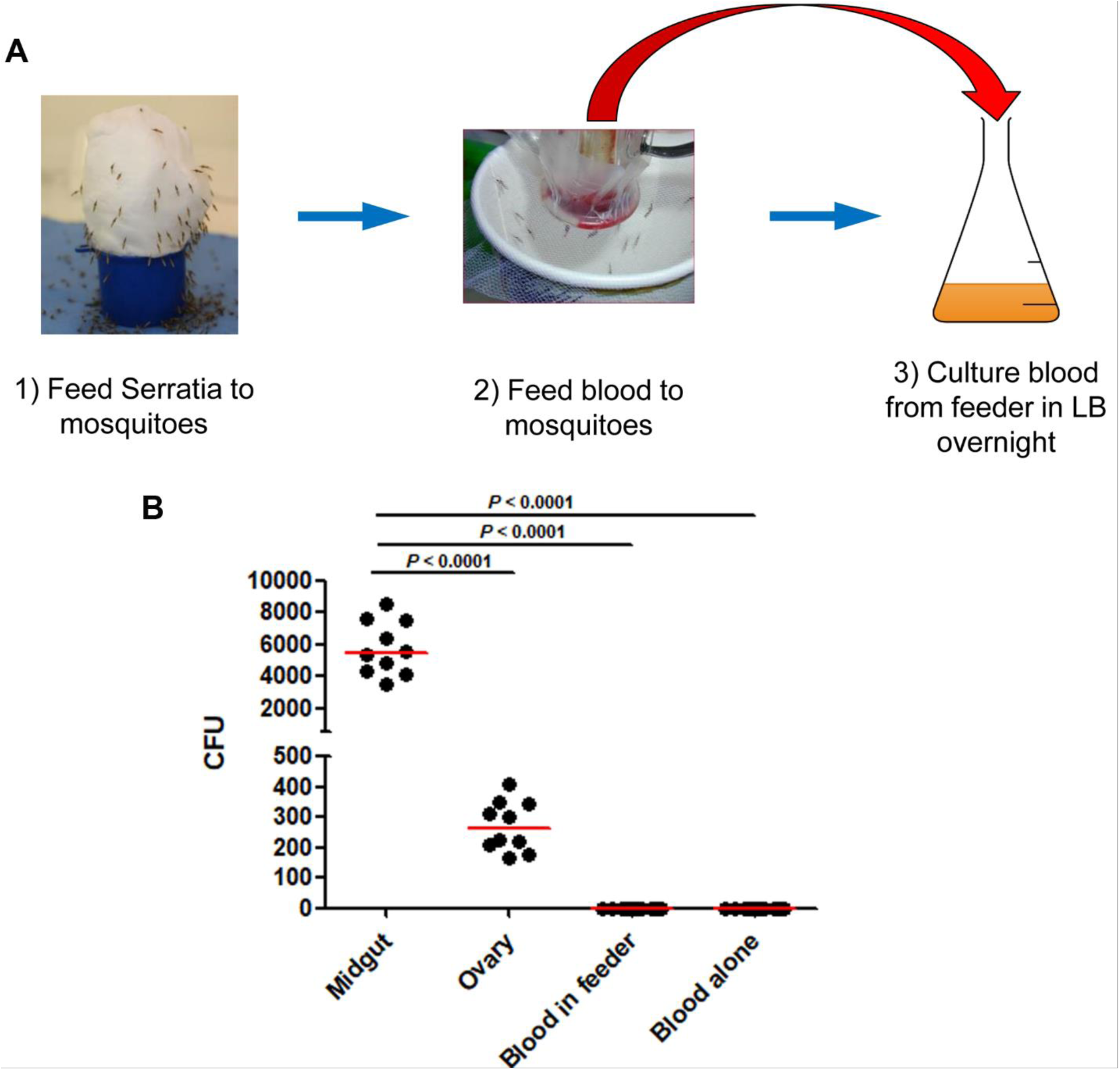
Schematic assay of bacteria transmitted from mosquitoes to the host using a membrane feeding assay. **[A]** Two-day-old mosquitoes were fed overnight on 10^7^ *Serratia*-GFP/ml of 5% sugar with food dye. Mosquitoes carrying the food dye marker were fed on sterile 5% sugar for two days. At this point, 100 female mosquitoes were starved for 3 h and fed blood for 30 min using a membrane feeder. To estimate the number of *Serratia*-GFP bacteria in the midgut and ovary, an additional 10 female mosquitoes were dissected prior to blood feeding, and dilutions of the homogenates were plated on LB/kanamycin plates. After blood feeding, 50 µl of blood from the feeder (out of 300 µl initial volume) was added to 5 ml LB/kanamycin, grown overnight, and then plated on LB/kanamycin plates to detect presence (or not) of *Serratia*-GFP in the blood. **[B]** Bacteria numbers in midguts, ovaries and blood from the feeder. No bacteria were detected from the blood samples. Data pooled from 10 independent biological experiments. Statistical analysis by the Mann–Whitney U test.

**Figure S4.**
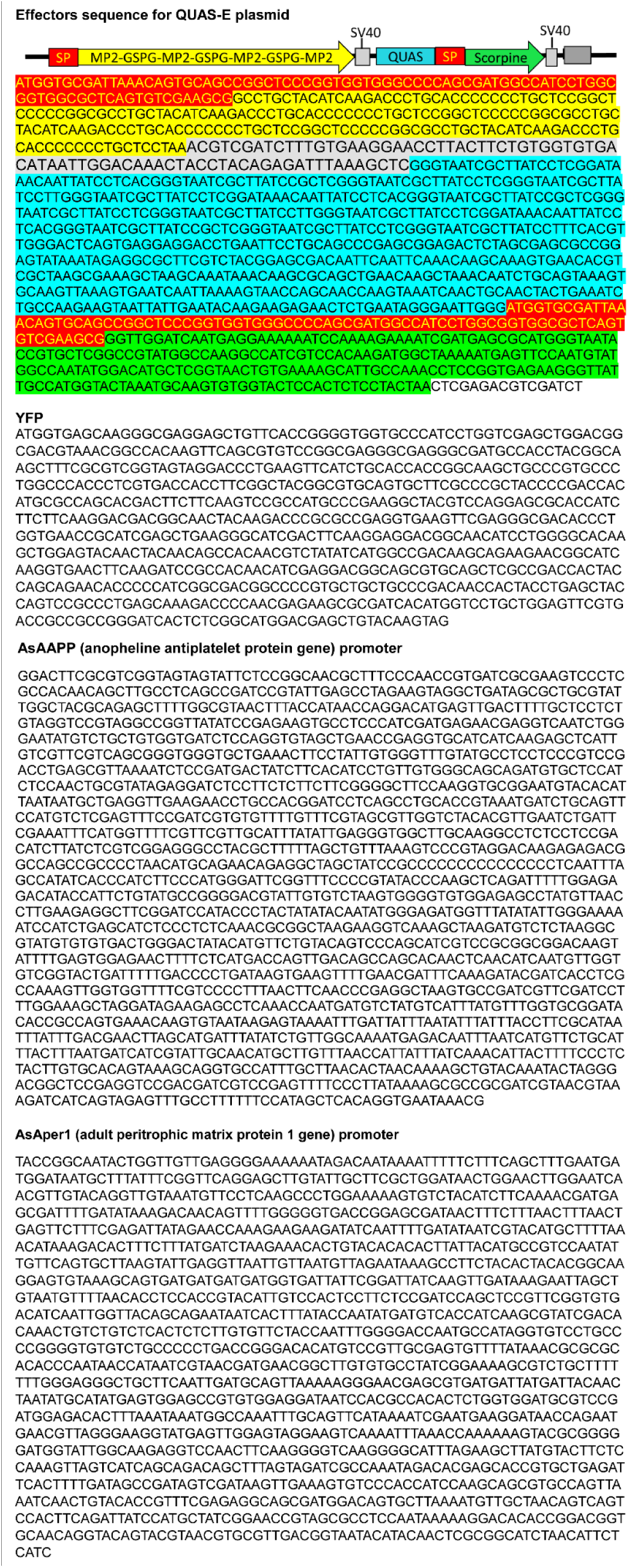
Sequences of the synthetic transgenes on the plasmid constructs for the transformation of *Anopheles* mosquito embryos.

**Table S1.**
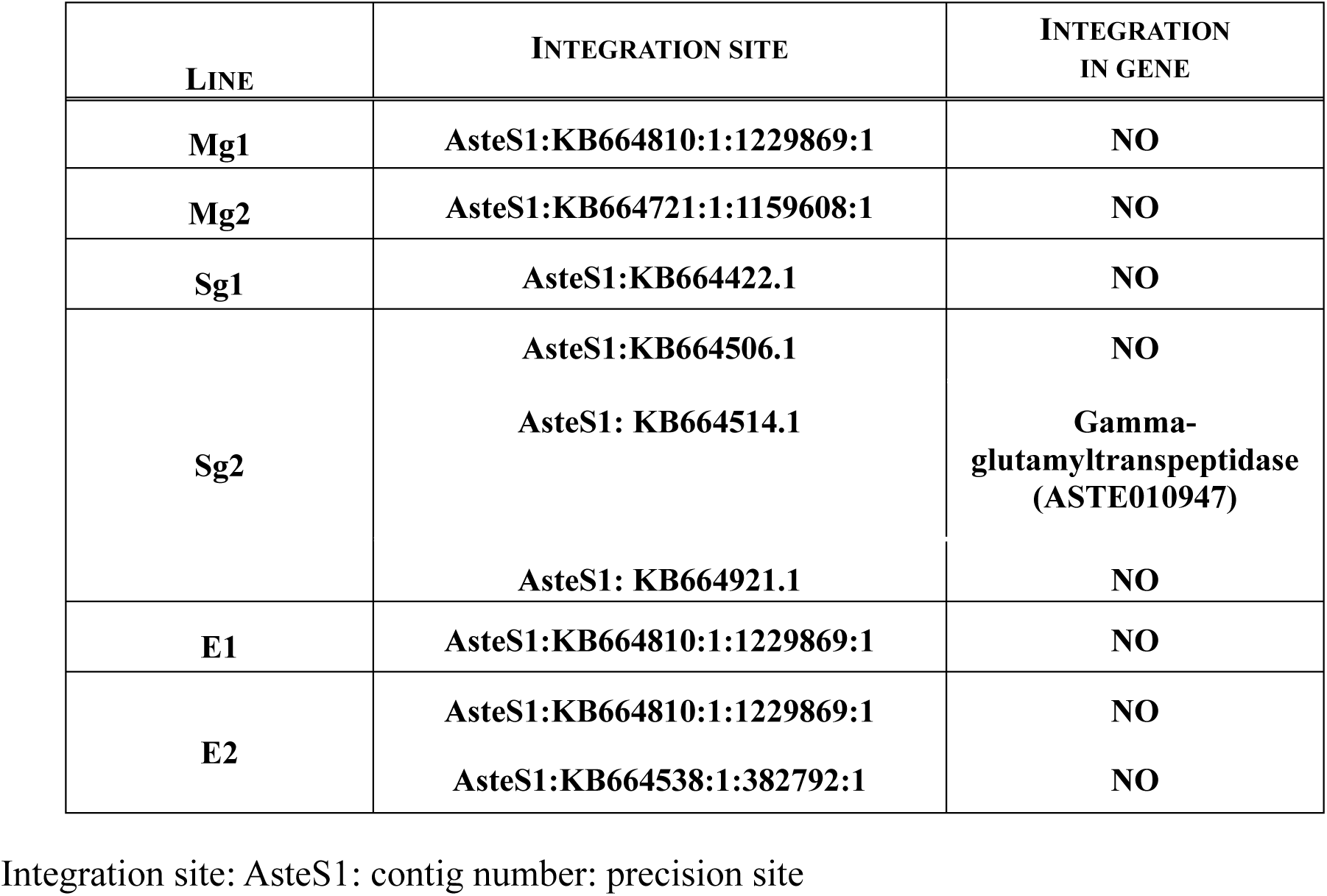
Transgene integration sites.

**Table S2.** Verification of transgene homozygosity.

| **MOSQUITO LINE** | **LARVA**<br>**NUMBERS** | **RED** | **YELLOW** | **BLUE** |
| --- | --- | --- | --- | --- |
| **Mg1** | **412** | **412** |  |  |
| **Mg2** | **276** | **276** |  |  |
| **Sg1** | **178** |  | **178** |  |
| **Sg2** | **329** |  | **329** |  |
| **E1** | **206** |  | **206** |  |
| **E2** | **378** |  | **378** |  |
| **Mg/E1** | **262** | **262** |  | **262** |
| **Mg/E2** | **198** | **198** |  | **198** |
| **Sg/E1** | **307** |  | **307** | **307** |
| **Sg/E2** | **345** |  | **345** | **345** |
| **Mg/Sg/E1** | **228** | **228** | **228** | **228** |
| **Mg/Sg/E2** | **361** | **361** | **361** | **361** |

A total of 20 transgenic female mosquitoes from each line were mated with wild type males and the progeny larvae assayed for expression of the dominant eye fluorescence marker. No not-fluorescent larvae were found, indicating that the females were homozygous for the transgenes.

**Table S3.** Expression of MP2 and scorpine mRNAs relative to the endogenous AsAper mRNA, quantified by qRT-PCR in the midgut of transgenic mosquitoes.

| Mosquito lines | Relative expression in midgut |  |  |
| --- | --- | --- | --- |
|  | AsAper | Scorpine | MP2 |
| E | 1.0± 0.1 | N | N |
| Mg/E | 1.0± 0.3 | 44.3±10.7 | 49.3±16.7 |
| Mg/Sg/E | 1.0± 0.2 | 56.1±8.7 | 35.6±8.9 |

**Table S4.**
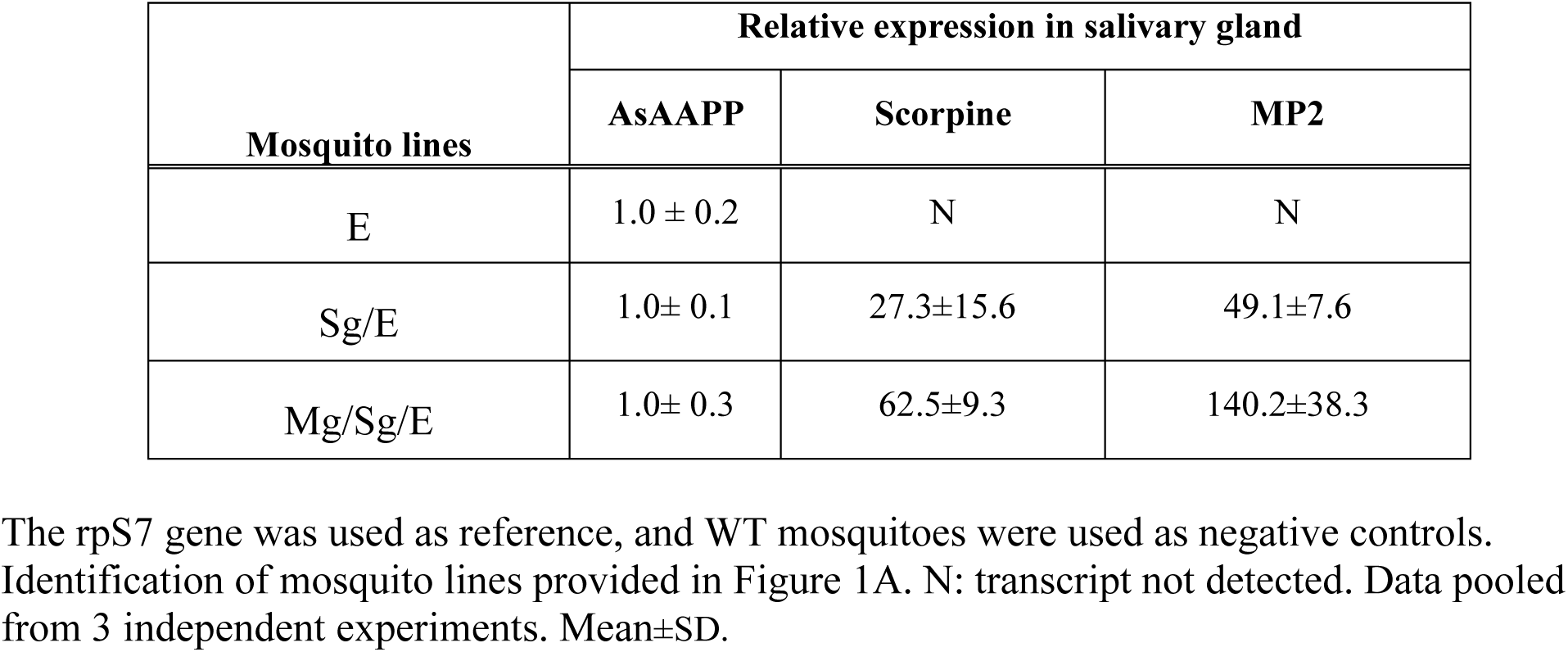
Relative expression of MP2 and scorpine mRNAs relative to the endogenous AsAAPP mRNA quantified by qRT-PCR in the salivary glands of transgenic mosquitoes.

| Mosquito lines | Relative expression in salivary gland |  |  |
| --- | --- | --- | --- |
|  | AsAAPP | Scorpine | MP2 |
| E | 1.0 ± 0.2 | N | N |
| Sg/E | 1.0± 0.1 | 27.3±15.6 | 49.1±7.6 |
| Mg/Sg/E | 1.0± 0.3 | 62.5±9.3 | 140.2±38.3 |

**Table S5.** *Serratia* is horizontally (sexually) transmitted.

| Males carrying<br>AS1-poly | Females<br>(mated/virgin) | Female CFUs |  |  |
| --- | --- | --- | --- | --- |
|  |  | Spermatheca | Midgut | Ovary |
| WT | WT mated | 0 | 0 | 0 |
| Transgenic | Transgenic mated | 0 | 0 | 0 |
| WT | WT virgin | 3.9±4.7 | 115±118 | 11±7.7 |
| Transgenic | Transgenic virgin | 2.9±4.7 | 104±111 | 8.7±7.8 |

Newly emerged virgin male adult mosquitoes were fed overnight on 5% sugar solution containing $10^7$ AS1-poly bacteria/ml and placed with females. Three days later, 10 females were assayed for the presence of *Serratia AS1* by plating midgut, ovary and spermatheca homogenates on apramycin/ampicillin agar plates and colonies were counted. Transgenic mosquito: Mg/Sg/E. Female mosquitoes mate only once in their lifetimes; mated females were used as controls. Data pooled from 3 independent experiments. Mean±SD.

**Table S6.** Vectors used in this research.

| **Vectors** | **Reference/notes** |
| --- | --- |
| phsp-pBac | 33 |
| pXL-BACII-DsRed-AAPP-QF2-hsp70 | 15 |
| pXL-BACIIIECFP-15XQUAS-TATA-PAI-SV40 | 15 |
| pXL-BACII- DsRed-Aper-QF2-Hsp70 | Mg QF2 driver plasmid |
| pXL-BAC-YFP-AAPP promoter-QF2-Hsp70 | Sg QF2 driver plasmid |
| pXL-BACIIIECFP-15XQUAS-TATA-MP2-SV40-15XQUAS-TATA-Scorpine-SV40 | QUAS-MP2-Scorpine effector plasmid |
| pBAM2-YFP | DNA template for YFP |

**Table S7.**
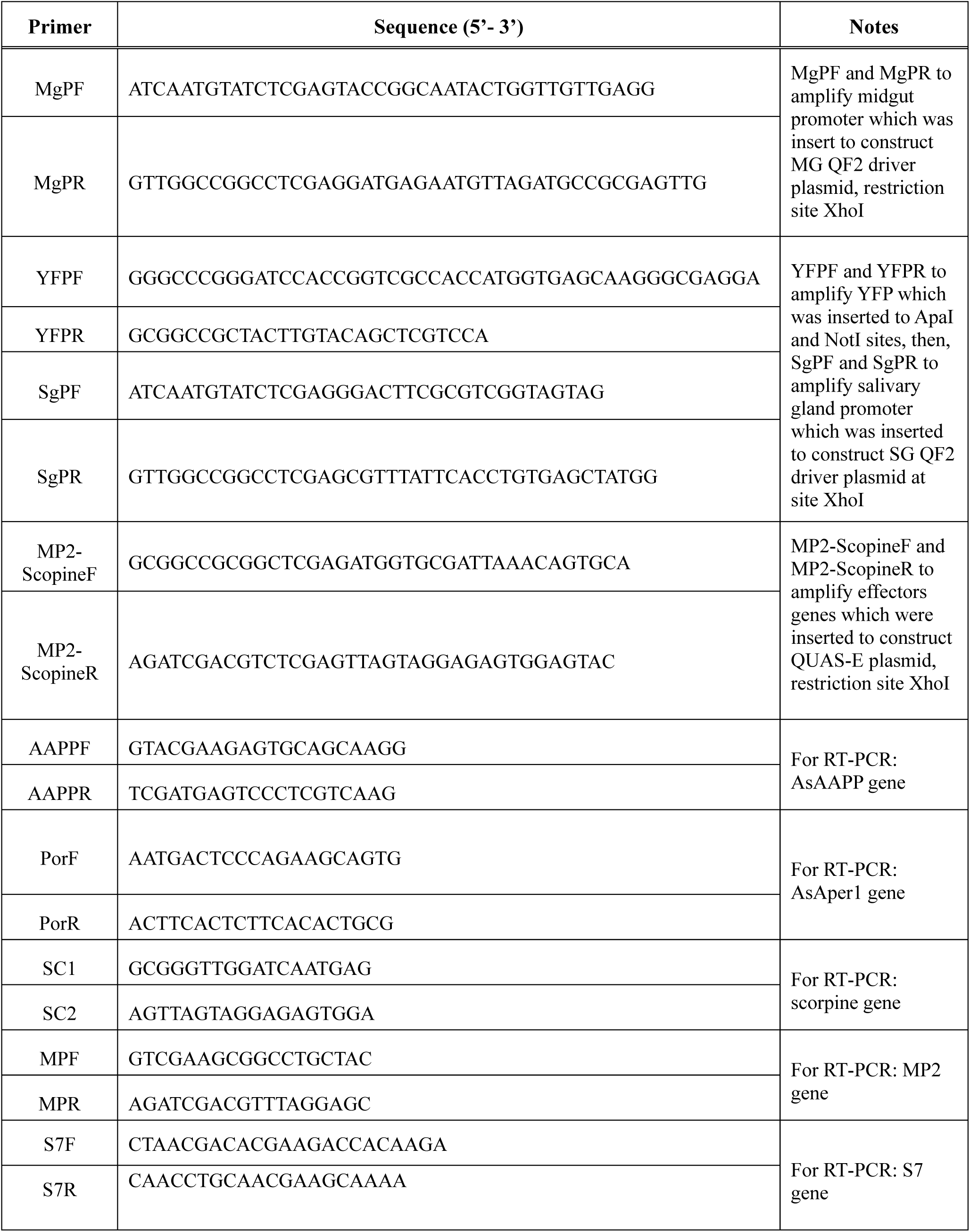
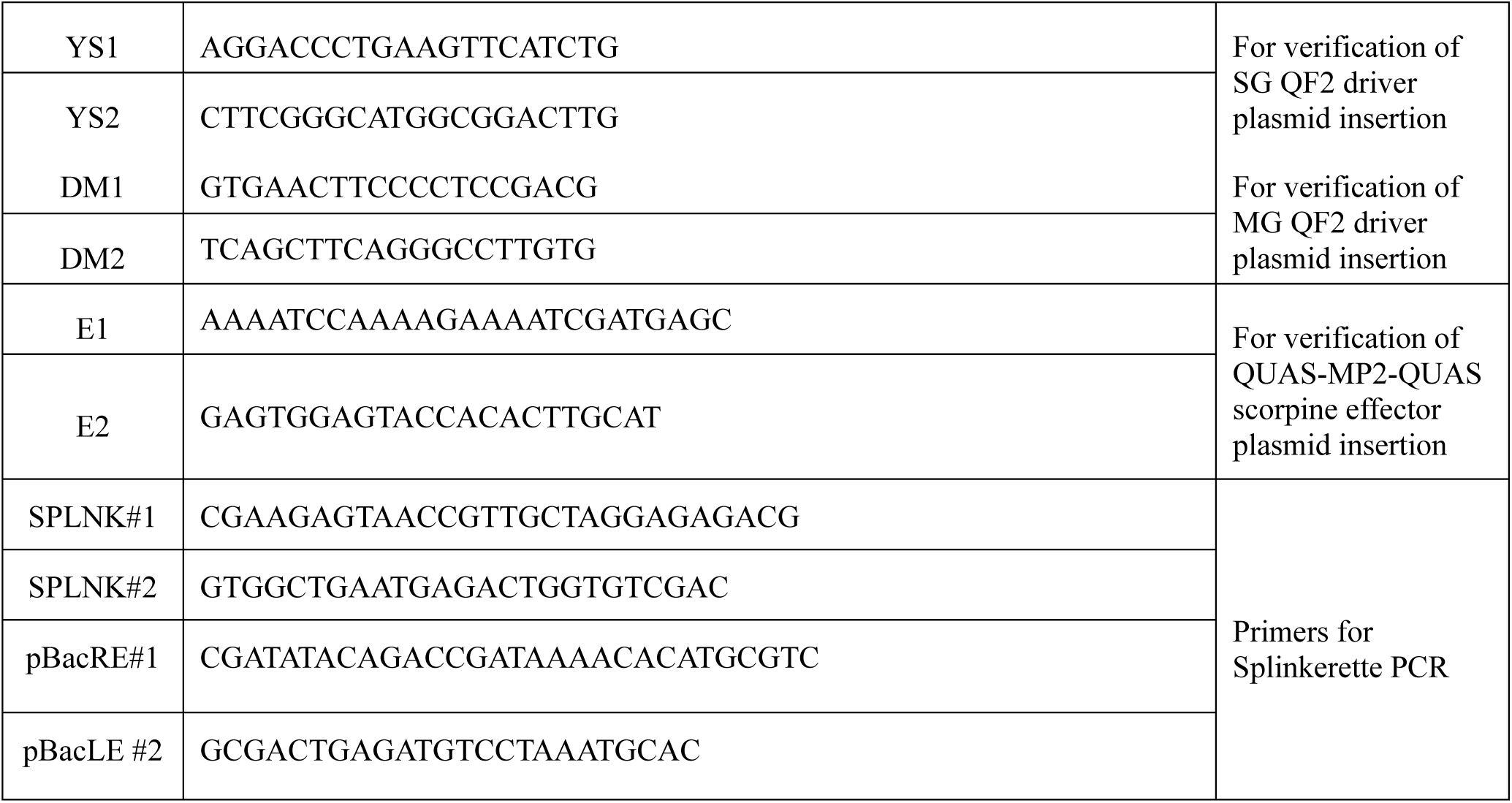
Oligonucleotide primers used in this study.

**Table S8.** Plasmid injection and screening for transformants.

| **Donor plasmid** | **Helper** | **# embryos injected** | **# G0 (number of survivors)** | **Pools** | **Pools with positive progeny** |
| --- | --- | --- | --- | --- | --- |
| MG QF2 driver plasmid | phsp-pBac | 440 | 35 | 17 | P1 and P2 |
| SG QF2 driver plasmid | phsp-pBac | 500 | 19 | 8 | P1 and P5 |
| QUAS-MP2-Scorpine effector plasmid | phsp-pBac | 547 | 43 | 22 | P1 and P2 |

## Notes

### Competing Interest Statement

The authors have declared no competing interest.

## References

1. World Health, O. (2020). World malaria report 2020: 20 years of global progress and challenges (World Health Organization).

2. Smith, R.C., Vega-Rodriguez, J., and Jacobs-Lorena, M. (2014). The Plasmodium bottleneck: malaria parasite losses in the mosquito vector. Mem Inst Oswaldo Cruz 109, 644–661.

3. Wang, S., and Jacobs-Lorena, M. (2013). Genetic approaches to interfere with malaria transmission by vector mosquitoes. Trends Biotechnol 31, 185–193. 10.1016/j.tibtech.2013.01.001.

4. Vanderberg, J.P. (1977). Plasmodium berghei: quantitation of sporozoites injected by mosquitoes feeding on a rodent host. Exp Parasitol 42, 169–181. 10.1016/0014-4894(77)90075-3.

5. Ito, J., Ghosh, A., Moreira, L.A., Wimmer, E.A., and Jacobs-Lorena, M. (2002). Transgenic anopheline mosquitoes impaired in transmission of a malaria parasite. Nature 417, 452–455. 10.1038/417452a.

6. Wang, S., Ghosh, A.K., Bongio, N., Stebbings, K.A., Lampe, D.J., and Jacobs-Lorena, M. (2012). Fighting malaria with engineered symbiotic bacteria from vector mosquitoes. Proc Natl Acad Sci U S A 109, 12734–12739. 10.1073/pnas.1204158109.

7. Dong, Y., Simoes, M.L., and Dimopoulos, G. (2020). Versatile transgenic multistage effector-gene combinations for Plasmodium falciparum suppression in Anopheles. Sci Adv 6, eaay5898. 10.1126/sciadv.aay5898.

8. Carballar-Lejarazu, R., Ogaugwu, C., Tushar, T., Kelsey, A., Pham, T.B., Murphy, J., Schmidt, H., Lee, Y., Lanzaro, G.C., and James, A.A. (2020). Next-generation gene drive for population modification of the malaria vector mosquito, Anopheles gambiae. Proc Natl Acad Sci U S A 117, 22805–22814. 10.1073/pnas.2010214117.

9. Quinn, C.M., and Nolan, T. (2020). Nuclease-based gene drives, an innovative tool for insect vector control: advantages and challenges of the technology. Curr Opin Insect Sci 39, 77–83. 10.1016/j.cois.2020.03.007.

10. Durvasula, R.V., Gumbs, A., Panackal, A., Kruglov, O., Aksoy, S., Merrifield, R.B., Richards, F.F., and Beard, C.B. (1997). Prevention of insect-borne disease: an approach using transgenic symbiotic bacteria. Proc Natl Acad Sci U S A 94, 3274–3278. 10.1073/pnas.94.7.3274.

11. Yoshida, S., Matsuoka, H., Luo, E., Iwai, K., Arai, M., Sinden, R.E., and Ishii, A. (1999). A single-chain antibody fragment specific for the Plasmodium berghei ookinete protein Pbs21 confers transmission blockade in the mosquito midgut. Mol Biochem Parasitol 104, 195–204. 10.1016/s0166-6851(99)00158-9.

12. Shane, J.L., Grogan, C.L., Cwalina, C., and Lampe, D.J. (2018). Blood meal-induced inhibition of vector-borne disease by transgenic microbiota. Nat Commun 9, 4127. 10.1038/s41467-018-06580-9.

13. Riehle, M.A., Moreira, C.K., Lampe, D., Lauzon, C., and Jacobs-Lorena, M. (2007). Using bacteria to express and display anti-Plasmodium molecules in the mosquito midgut. Int J Parasitol 37, 595–603. 10.1016/j.ijpara.2006.12.002.

14. Wang, S., Dos-Santos, A.L.A., Huang, W., Liu, K.C., Oshaghi, M.A., Wei, G., Agre, P., and Jacobs-Lorena, M. (2017). Driving mosquito refractoriness to Plasmodium falciparum with engineered symbiotic bacteria. Science 357, 1399–1402. 10.1126/science.aan5478.

15. Potter, C.J., Tasic, B., Russler, E.V., Liang, L., and Luo, L. (2010). The Q system: a repressible binary system for transgene expression, lineage tracing, and mosaic analysis. Cell 141, 536–548. 10.1016/j.cell.2010.02.025.

16. Riabinina, O., Task, D., Marr, E., Lin, C.C., Alford, R., O’Brochta, D.A., and Potter, C.J. (2016). Organization of olfactory centres in the malaria mosquito Anopheles gambiae. Nat Commun 7, 13010. 10.1038/ncomms13010.

17. Shen, Z., and Jacobs-Lorena, M. (1998). A type I peritrophic matrix protein from the malaria vector Anopheles gambiae binds to chitin. Cloning, expression, and characterization. J Biol Chem 273, 17665–17670. 10.1074/jbc.273.28.17665.

18. Abraham, E.G., Donnelly-Doman, M., Fujioka, H., Ghosh, A., Moreira, L., and Jacobs-Lorena, M. (2005). Driving midgut-specific expression and secretion of a foreign protein in transgenic mosquitoes with AgAper1 regulatory elements. Insect Mol Biol 14, 271–279. 10.1111/j.1365-2583.2004.00557.x.

19. Yoshida, S., and Watanabe, H. (2006). Robust salivary gland-specific transgene expression in Anopheles stephensi mosquito. Insect Mol Biol 15, 403–410. 10.1111/j.1365-2583.2006.00645.x.

20. Vega-Rodriguez, J., Ghosh, A.K., Kanzok, S.M., Dinglasan, R.R., Wang, S., Bongio, N.J., Kalume, D.E., Miura, K., Long, C.A., Pandey, A., and Jacobs-Lorena, M. (2014). Multiple pathways for Plasmodium ookinete invasion of the mosquito midgut. Proc Natl Acad Sci U S A 111, E492–500. 10.1073/pnas.1315517111.

21. Conde, R., Zamudio, F.Z., Rodriguez, M.H., and Possani, L.D. (2000). Scorpine, an anti-malaria and anti-bacterial agent purified from scorpion venom. FEBS Lett 471, 165–168. 10.1016/s0014-5793(00)01384-3.

22. Gao, B., Xu, J., Rodriguez Mdel, C., Lanz-Mendoza, H., Hernandez-Rivas, R., Du, W., and Zhu, S. (2010). Characterization of two linear cationic antimalarial peptides in the scorpion Mesobuthus eupeus. Biochimie 92, 350–359. 10.1016/j.biochi.2010.01.011.

23. Potter, C.J., and Luo, L. (2010). Splinkerette PCR for mapping transposable elements in Drosophila. PLoS One 5, e10168. 10.1371/journal.pone.0010168.

24. Fang, W., Vega-Rodriguez, J., Ghosh, A.K., Jacobs-Lorena, M., Kang, A., and St Leger, R.J. (2011). Development of transgenic fungi that kill human malaria parasites in mosquitoes. Science 331, 1074–1077. 10.1126/science.1199115.

25. Luo, E., Matsuoka, H., Yoshida, S., Iwai, K., Arai, M., and Ishii, A. (2000). Changes in salivary proteins during feeding and detection of salivary proteins in the midgut after feeding in a malaria vector mosquito, Anopheles stephensi (Diptera : Culicidae). Medical Entomology and Zoology 51, 13–20. 10.7601/mez.51.13.

26. Carballar-Lejarazu, R., Rodriguez, M.H., de la Cruz Hernandez-Hernandez, F., Ramos-Castaneda, J., Possani, L.D., Zurita-Ortega, M., Reynaud-Garza, E., Hernandez-Rivas, R., Loukeris, T., Lycett, G., and Lanz-Mendoza, H. (2008). Recombinant scorpine: a multifunctional antimicrobial peptide with activity against different pathogens. Cell Mol Life Sci 65, 3081–3092. 10.1007/s00018-008-8250-8.

27. Nolan, T. (2021). Control of malaria-transmitting mosquitoes using gene drives. Philos Trans R Soc Lond B Biol Sci 376, 20190803. 10.1098/rstb.2019.0803.

28. Simoni, A., Hammond, A.M., Beaghton, A.K., Galizi, R., Taxiarchi, C., Kyrou, K., Meacci, D., Gribble, M., Morselli, G., Burt, A., et al. (2020). A male-biased sex-distorter gene drive for the human malaria vector Anopheles gambiae. Nat Biotechnol 38, 1054–1060. 10.1038/s41587-020-0508-1.

29. Adolfi, A., Gantz, V.M., Jasinskiene, N., Lee, H.F., Hwang, K., Terradas, G., Bulger, E.A., Ramaiah, A., Bennett, J.B., Emerson, J.J., et al. (2020). Efficient population modification gene-drive rescue system in the malaria mosquito Anopheles stephensi. Nat Commun 11, 5553. 10.1038/s41467-020-19426-0.

30. Gao, H., Bai, L., Jiang, Y., Huang, W., Wang, L., Li, S., Zhu, G., Wang, D., Huang, Z., Li, X., et al. (2021). A natural symbiotic bacterium drives mosquito refractoriness to Plasmodium infection via secretion of an antimalarial lipase. Nat Microbiol 6, 806–817. 10.1038/s41564-021-00899-8.

31. Bilgo, E., Vantaux, A., Sanon, A., Ilboudo, S., Dabire, R.K., Jacobs-Lorena, M., and Diabate, A. (2018). Field assessment of potential sugar feeding stations for disseminating bacteria in a paratransgenic approach to control malaria. Malar J 17, 367. 10.1186/s12936-018-2516-x.

32. Feldmann, A.M., and Ponnudurai, T. (1989). Selection of Anopheles stephensi for refractoriness and susceptibility to Plasmodium falciparum. Med Vet Entomol 3, 41–52. 10.1111/j.1365-2915.1989.tb00473.x.

33. Huang, W., Wang, S., and Jacobs-Lorena, M. (2020). Self-limiting paratransgenesis. PLoS Negl Trop Dis 14, e0008542. 10.1371/journal.pntd.0008542.

34. Tripathi, A.K., Mlambo, G., Kanatani, S., Sinnis, P., and Dimopoulos, G. (2020). Plasmodium falciparum Gametocyte Culture and Mosquito Infection Through Artificial Membrane Feeding. J Vis Exp. 10.3791/61426.

35. Volohonsky, G., Terenzi, O., Soichot, J., Naujoks, D.A., Nolan, T., Windbichler, N., Kapps, D., Smidler, A.L., Vittu, A., Costa, G., et al. (2015). Tools for Anopheles gambiae Transgenesis. G3 (Bethesda) 5, 1151–1163. 10.1534/g3.115.016808.

36. Zhang, J., Huang, F.S., Xu, W.Y., Wang, Y., Zhou, T.L., and Duan, J.H. (2011). Plasmodium yoelii: correlation of TEP1 with mosquito melanization induced by nitroquine. Exp Parasitol 127, 52–57. 10.1016/j.exppara.2010.06.032.

37. Cha, S.J., McLean, K.J., and Jacobs-Lorena, M. (2018). Identification of Plasmodium GAPDH epitopes for generation of antibodies that inhibit malaria infection. Life Sci Alliance 1, e201800111. 10.26508/lsa.201800111.

38. Alves, E.S.T.L., Radtke, A., Balaban, A., Pascini, T.V., Pala, Z.R., Roth, A., Alvarenga, P.H., Jeong, Y.J., Olivas, J., Ghosh, A.K., et al. (2021). The fibrinolytic system enables the onset of Plasmodium infection in the mosquito vector and the mammalian host. Sci Adv 7. 10.1126/sciadv.abe3362.

